# Statistical mechanics of phenotypic eco-evolution: from adaptive dynamics to complex diversification

**DOI:** 10.1101/2023.07.20.549856

**Authors:** Matteo Sireci, Miguel A. Muñoz

## Abstract

The ecological and evolutionary dynamics of large sets of individuals can be theoretically addressed using ideas and tools from statistical mechanics. This strategy has been addressed in the literature, both in the context of population genetics –whose focus is of genes or “genotypes”— and in adaptive dynamics, putting the emphasis on traits or “phenotypes”. Following this tradition, here we construct a framework allowing us to derive “macroscopic” evolutionary equations from a rather general “microscopic” stochastic dynamics representing the fundamental processes of reproduction, mutation and selection in a large community of individuals, each one characterized by its phenotypic features. Importantly, in our setup, ecological and evolutionary timescales are intertwined, which makes it particularly suitable to describe microbial communities, a timely topic of utmost relevance. Our framework leads to a probabilistic description of the distribution of individuals in phenotypic space —even in the case of arbitrarily large populations— as encoded in what we call “generalized Crow-Kimura equation” or “generalized replicator-mutator equation”. We discuss the limits in which such an equation reduces to the (deterministic) theory of “adaptive dynamics” (i.e. the standard approach to evolutionary dynamics in phenotypic space. Moreover, we emphasize the aspects of the theory that are beyond the reach of standard adaptive dynamics. In particular, by working out, as a guiding example, a simple model of a growing and competing population, we show that the resulting probability distribution can exhibit “dynamical phase transitions” changing from unimodal to bimodal —by means of an evolutionary branching— or to multimodal, in a cascade of evolutionary branching events. Furthermore, our formalism allows us to rationalize these cascades of transitions using the parsimonious approach of Landau’s theory of phase transitions. Finally, we extend the theory to account for finite populations and illustrate the possible consequences of the resulting stochastic or “demographic” effects. Altogether the present framework extends and/or complements existing approaches to evolutionary/adaptive dynamics and paves the way to more systematic studies of e.g. microbial communities as well as to future developments including theoretical analyses of the evolutionary process from the general perspective of non-equilibrium statistical mechanics.

## I. INTRODUCTION

Darwinian evolution relies on the fundamental principles of reproduction, mutation and selection, to describe how populations change in time and how new forms branch out from existing ones [1, 2]. The theory of evolutionary dynamics —whose aim is that of formalizing the ideas of Darwinian evolution from a conceptual and quantitative perspective— has become a wide and mature discipline at the crossroad between biology, mathematics and statistical physics [3–9]. Diverse theoretical approaches, differing in various aspects, have been proposed to model and rationalize evolutionary phenomena [10] but, rather generically, evolutionary dynamics is formulated building on the theories of dynamical-systems and stochastic processes [11–16].

In spite of the broadness of this territory, some overarching concepts and principles have been put forward. For instance, Price in his seminal paper “*Mathematical Theory of Selection*”, set the bases for the development of a general and abstract mathematical theory of selection [17]. Actually, the so-called “Price equation” quantifies how the abundance of a given gene or phenotypic trait changes as a function of its covariance with its associated fitness [17–24] and is, hence, sometimes termed the “algebra of evolution” [14, 19, 21]. Later, Page and Nowak showed that different *“macroscopic”* formulations of evolutionary dynamics may actually be equivalent to each other [10, 25–27], even if specific details depend on whether the focus is on genes (*genotypic evolution*) or on phenotypic traits (*phenotypic evolution*).

*Population genetics* —i.e. the study of genetic variations within a population (or between different populations) together with the evolutionary factors causing such a variation— was developed starting from classical works of Fisher [28, 29], Wright [30], Price [18], Kimura [11, 12] and others, including recent exciting developments [31–35]. On the other hand, quantitative approaches to evolutionary dynamics focusing on phenotypes or traits (rather than on “genotypes”) were developed later, in the last decades. These include *evolutionary game theory* [15, 36–42], which aims to study possible equilibria of different populations with discrete and fixed traits (or “strategies”) as well as *adaptive dynamics* [43–52], which includes the possibility of mutations, so that phenotypes are not discrete but can change in a continuum (see below).

A common assumption often considered in these theoretical approaches is that mutations occur at a relatively low pace so that ecological (fast) and evolutionary (slow) timescales are well separated [53, 54]. This assumption which has been historically retained as a very natural one is not, however, universally valid, and may fail dramatically, especially when dealing with populations of micro-organisms (see, e.g., [55–58]).

Microbial communities are the most abundant and diverse ones in the biosphere [59, 60]. Studying their dynamics has become a crucial challenge from many diverse viewpoints including, e.g., environmental and health-related perspectives [61, 62]. As a matter of fact, microbial communities have become the ultimate frontier and test bench of modern evolutionary theory [63]. Indeed, microbial communities often evolve very rapidly, with frequent mutations and fast selection; for instance, viruses and bacteria may have astonishingly-high mutation rates [64–67]. Consequently, evolutionary effects cannot be decoupled from ecological ones, as both can occur at contemporary timescales [68, 69] and microbial communities often exhibit a very large fine-scale diversity in the form of multiple co-occurring phenotypes [70]. Importantly, such a diversity is nowadays accessible to experiments owing to technological advances in determining single-cell traits [31, 71, 72] and metabolic functions [73–76]. All this calls for the development of novel eco-evolutionary frameworks —extending existing ones— to analyze complex microbial communities, with individuals distributed in phenotypic space and evolving on ecological time scales.

The standard theory of phenotypic evolution is *adaptive dynamics* [43–52]. In adaptive dynamics (AD) a population is assumed to be in a steady state, called “resident type”, and small stochastic variations of such a type, i.e. “mutants”, are assumed to emerge at a very slow rate. In the case that the per-capita growth rate of the mutant within the resident-type population —i.e, its “invasion fitness”— is positive, the mutant is assumed to invade the population and, eventually, become fixated as the new resident type [77].

Within the standard AD framework, mutations are typically considered to be: (i) rare (which implicitly assumes a large separation of timescales between ecological and evolutionary processes), such that the system has time to re-equilibrate to a new steady state after each mutation; (ii) small (as a result of which the probability distributions in phenotypic space are typically Gaussians); (iii) independent of the parental traits; and (iv) not subject themselves to evolution [43, 45, 48, 51, 78].

Since its original formulation, AD enjoyed a great success, as it allowed to unify evolutionary dynamics with realistic ecological features [51]. For instance, remarkably, AD allows for the possibility to account for “*evolutionary branching*” –i.e. the split of an initially monomorphic population into two diverse sub-populations— shedding light on how speciation [44, 45, 48, 50, 51, 79, 80] and diversification in sympatry [78] may come about. Similarly, phenomena such as the evolution of dispersal strategies [81], pathogenicity [82], metabolic preferences [83, 84], cancer [85, 86] multi-cellularity [87], to name but a few, have been successfully addressed within the context of AD. Moreover, extensions of AD have been developed to include ingredients such as finite populations [47, 88], species interactions [27], sexual populations [48], multi-dimensional phenotypic spaces [29, 89–92], variable environmental conditions [93], or variability in the evolutionary outcomes [94]. Nevertheless, given the above-mentioned restrictive hypotheses, there remains space to generalize AD in a number of directions.

From the perspective of statistical mechanics, it would be highly desirable to construct a general stochastic individual-based (“many-body” or “many-particle”) theory for agents in a community exposed to basic rules of reproduction, mutation, and selection, such that it could reproduce deterministic theories such as AD in some large-system-size limit. The goal would be to be able to derive “macroscopic” probabilistic equations for the evolution of populations and communities —starting from a “microscopic” description of stochastic processes acting at the level of interacting individuals (including ecological and evolutionary processes at comparable timescales)—by employing the powerful methods of statistical physics.

Before proceeding toward such an ambitious goal, let us remark that diverse approaches have already tackled the previous challenge and that significant advances have already been made in this direction. In particular, we extensively elaborate and rely —among others— upon the following works:

- Dieckmann and Law were pioneers in deriving the deterministic equations of AD using a probabilistic description of the population in phenotypic space. Their approach allows e.g. for the possibility that potentially-successful mutants become accidentally extinct owing to demographic fluctuations before achieving fixation [47].
- Champagnat and colleagues built a rather rigorous mathematical framework, allowing them to derive the “macroscopic” equations of AD starting from “microscopic” underlying birth-and-death stochastic processes and generalized AD in various ways [95–98].
- Frey and coauthors developed a formalism to derive a macroscopic equation from the underlying microscopic birth-death process in the context of bacterial populations [99, 100].
- Wakano and Iwasa [88] studied the effects of demographic fluctuations within the context of AD (see also [80, 98]).

In a similar spirit to these and related approaches, our aim in what follows is to develop a general framework deeply rooted in the views and methods of statistical mechanics, able to generalize AD to eco-evolutionary scenarios. In particular, we present a probabilistic theory of the evolution of *trait distributions* [101] including the effect of selection, arbitrary mutations, and fluctuations stemming from finite population sizes, where ecological and evolutionary processes occur contemporarily. [102]

The present work contributes to the development of an eco-evolutionary theory for microbial communities, allowing to shed further light on the empirically-observed astonishing diversity in traits and interactions of microbial communities. Our hope is that it makes this kind of quantitative approaches to complex eco-evolutionary communities accessible to a broader audience, including physicists, biologists, and ecologists.

The manuscript is organized as follows. In Sec.II, we introduce our general eco-evolutionary framework. Starting with a rather general (*microscopic*) individual-based birth-death process involving reproduction, selection, and mutation (Sec.II A), we derive a *macroscopic* equation describing the evolution of a general trait distribution in phenotypic space (Sec.II B). From this, we derive equations for its moments (Sec.II C) and particularize them to the case of small mutations (Sec.II D). This allows us to recover the standard theory of AD using a Gaussian approximation for the trait distribution (Sec.III) as well as to formulate an extended theory “*a la Landau*”, including higher moments in the expansion, to go beyond AD (Sec.IV). The general theory is then applied to the specific example of individual-based model including both a fixed external ecological niche and competition among individuals allowing us to illustrate its richness. Finally, we extend our deterministic theory to account for demographic stochastic effects for finite populations (Sec.VI). To close, we discuss our main conclusions as well as the limitations of our approach and possible future developments (Sec.VII).

## II. GENERAL FRAMEWORK AND MACROSCOPIC ECO-EVOLUTIONARY EQUATIONS

For the sake of simplicity and without loss of generality, we consider —as customarily done in evolutionary dynamics [103]— a population of fixed size, composed of *N* individuals. As in AD, we choose to focus on a phenotypic description of individuals. Thus, each of them (say the *i*-th one, with *i*∈ [1, *N*]) is characterized by a set of *phenotypic traits* that, in the simplest possible case, can be encoded in a single real-valued variable, *x*_*i*_. This represents a coordinate in a one-dimensional phenotypic space 𝒫. Generalizations to higher-dimensional phenotypic spaces and to populations of variable size can easily be addressed.

The population as a whole can be represented by a *N* -dimensional vector ***x*** = (*x*_1_, *x*_2_, *x*_3_, .., *x*_*N*_), that we call a *phenotypic configuration*. The final goal is to describe the dynamics of the probability *P* (***x***, *t*) to have a population with a given phenotypic configuration ***x*** as a function of time.

### A. Birth-and-death eco-evolutionary model

The dynamics of *P* (***x***, *t*) can be mathematically described by a master equation, which encodes the main stochastic processes occurring at an individual or “microscopic” level [104]. In particular, the stochastic dynamics at the individual level relies on the three key ingredients of Darwinian evolution [1, 14] (see the sketch of Fig. 1):

**FIG. 1.**
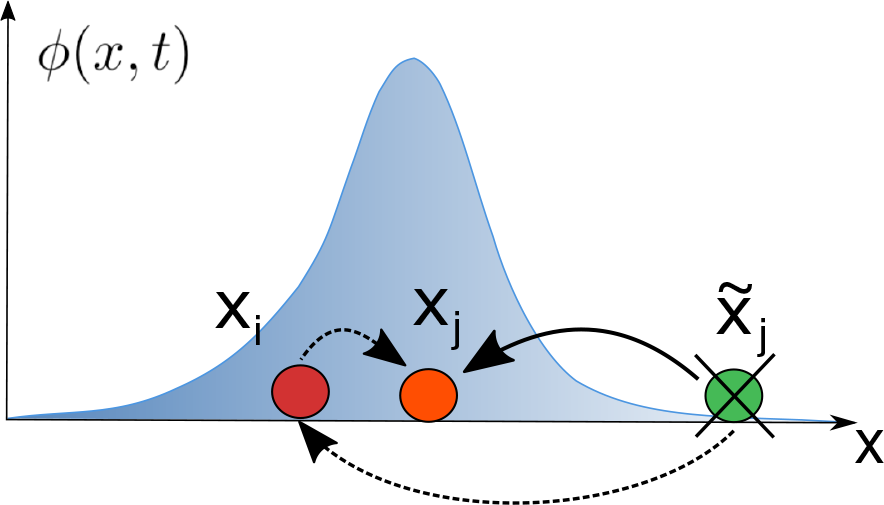
Sketch of the basic processes characterizing the eco-evolutionary dynamics at an individual (“microscopic”) level. The probability density *ϕ*(*x, t*) describes the fraction of the population with phenotypic trait *x* at a given time *t*. This can change as a result of the following stochastic processes: (i) *reproduction with mutation* (individual with trait *x*_*i*_ (red) generates an offspring with trait *x*_*j*_ relatively close —but not identical— to *x*_*i*_ (orange); (ii) *death* (an individual with trait 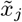(green) is randomly selected to die. Mathematically, these processes are described by a Markov process (as represented by black dashed arrows) in which the individual *i* performs a *local jump* to the coordinate *x*_*j*_, while its initial position *x*_*i*_ becomes occupied by the removed individual by means of a *non-local* jump from 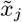 . Moreover, the composition of these two transitions can be effectively described as a single net jump (solid black line) from 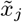 to *x*_*j*_ . These processes give rise to an evolving distribution that may eventually converge to a stationary state.

#### (i) Reproduction

Each individual *i* produces (asexually) one offspring at some transition rate, called “fitness”, *f*_*i*_(***x***) *>* 0, that depends on its phenotype as well as on the overall system’s state ***x***. In what follows, we restrict ourselves to the case in which the fitness for individuals with trait *x*_*i*_ is composed of (i) an intrinsic growth rate *K*(*x*_*i*_) —that is assumed to depend only on the individual trait *x*_*i*_— and defines an external “ecological niche”, specific of the considered environment/conditions, as well as (ii) an additional term that comes from pairwise ecological forces (or “interactions”), *I*(*x*_*i*_, *x*_*j*_) which represent competition, cooperation, and/or more complex ecological forces with all other individuals (e.g. the *j*-th one with trait value *x*_*j*_). Thus, the total fitness function *f*_*i*_(***x***) of individual *i* is given by the intrinsic growth rate plus the weighted sum of all the pairwise interactions with other individuals (so that both terms are typically of the same order)

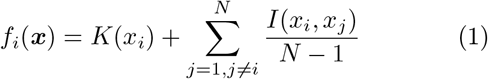

or, equivalently

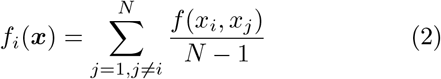

Where

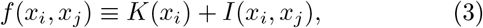

includes up to pairwise interactions. Let us stress that higher-order interactions, which might be relevant in some cases [105], could also be straightforwardly implemented by including additional terms, depending on more than two individual traits, in Eq.(1)).

#### (ii) Selection

The fact that individuals with larger fitness values are more likely to reproduce allows one to implement natural selection in an indirect way just by imposing a constant population size, *N* . In particular, as illustrated in Fig. 1, our model assumes that each time that an individual *i* produces an offspring, a second individual *j* (different from *i*) is randomly chosen and removed from the community [106]. This second individual is selected with certain death probability *d*_*j*_(***x***) which, in general can depend on the full phenotypic state, ***x***, and that, for simplicity, we set to be constant across time and individuals, i.e. *d*_*j*_ = 1*/*(*N* − 1).

#### (iii) Heredity and variation

When an individual replicates, its offspring inherits the phenotypic trait/s of the parent, with some variation. The probability of a given mutation from the mother value *x*_*i*_ to the offspring’s one *x*_*j*_ is represented by a generic distribution function, *β*(*x*_*i*_, *x*_*j*_), called “*mutation kernel* “. This distribution can be characterized by its mean *θ* and variance *σ*, which in simple cases (but not always) can be assumed to be independent of *x*_*i*_ and depend only on the magnitude of the jump *δ* = | *x*_*i*_ −*x*_*j*_| . Otherwise, when the jumps are state dependent the function can be parametrized as *β*(*x, δ*).

Thus, the master equation defining the general model (see Fig. 1 for a sketch) can be written as [104]:

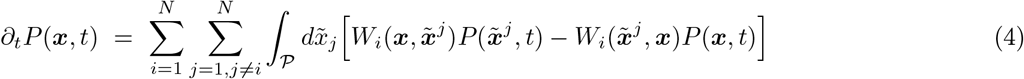

Where

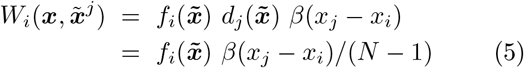

is the rate to transition from an initial state 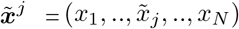 to ***x*** = (*x*_1_, .., *x*_*j*_, .., *x*_*N*_), where 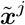 differs from ***x*** only in the value of the coordinate *j*, i.e. the individual that is killed and replaced by an offspring of *i* with mutated trait *x*_*j*_. Reciprocally, 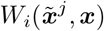 is the transition rate for the reverse process.

Let us remark that the described stochastic dynamics generates a neat flux of probability from the phenotypic state of the removed individual (i.e. trait 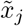) to that of the newly generated one, (i.e. *x*_*j*_), which can effectively be visualized as a direct jump from the first position to the latter (solid line in Fig. 1), though it actually consists of two different jumps (dashed lines in Figure 1): a non-local one, from 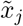 to *x*_*i*_, and a local one from the latter to its offspring *x*_*j*_. The first (non-local) jump implements the effect of selection while the second (local) one describes mutation.

### B. Selection-mutation mean-field equation

The above master equation, Eq. (4), is very general but, clearly, difficult to handle analytically. Thus, in order to gain more insight, here we employ some standard approximations —that, later on, will be relaxed— to reduce it to a simpler deterministic (mean-field like) equation for the marginal probability to find any individual in a particular state *x, ϕ*(*x, t*).

Readers not interested in the formal aspects of the following mathematical derivations can go directly to Eq.(10).

As often done in statistical mechanics, one can assume that individuals are, *a priori*, indistinguishable. This equivalence allows one to derive an equation for the individual (or “one-particle”) probability density distribution *ϕ*(*x, t*) from Eq. (4). More specifically: the *density* of individuals with phenotype *x* at time *t* in a given realization of the stochastic process can be simply expressed as

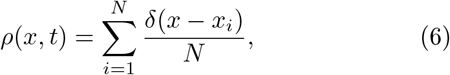

and averaging over many possible noise realizations one can derive the probability density distribution

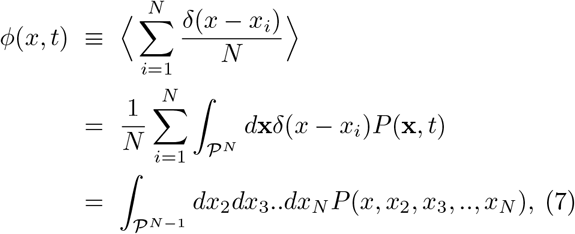

that is nothing but the marginal probability of one of the individuals, with phenotype (e.g.) *x* ≡ *x*_1_, given that the probability distribution is symmetric with respect to the exchange of individual labels [99, 107].

Taking the time derivative of Eq.(7) and using the master equation, Eq.(4), one readily obtains (see SI, Sec B.):

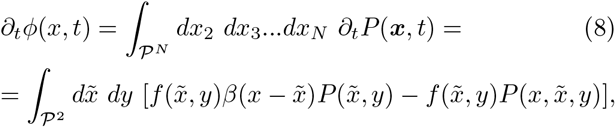

where 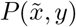 and 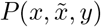 are 2-particle and 3-particle joint probabilities.

Before proceeding, let us remark that there are two main consequences of individual indistinguishability:

1. The fitness function has been reduced to the pairwise fitness function, *f* (*x, y*); i.e. it depends just on the traits of the reproducing individual, x, and another *generic one*, y, —specified only by its trait— with which it interacts.
2. The resulting simplified master equation, Eq.(8), depends only on the two- and three-particle joint probability (rather than in the whole N-particle one) so that it is not a closed equation for the 1-particle density, *ϕ*(*x, t*). In order to obtain a closed equation for *ϕ*(*x, t*) one needs to make the additional the mean-field approximation consisting, as usual, in assuming that the joint probability distribution function can be factorized:

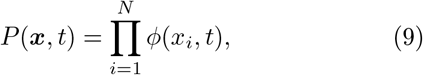

which is an exact result in the limit of infinitely large population sizes [99, 100, 107]. Corrections accounting for finite population size will be explicitly introduced and discussed in a forthcoming section (see Sec.VI).

This readily leads to the deterministic (or mean-field) equation:

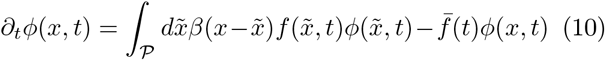

where we have defined the “*marginal fitness*” associated with state 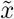

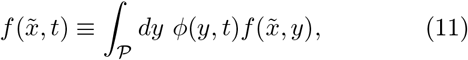

with 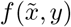 defined by Eq.(3) and where 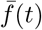 is the population-averaged marginal fitness:

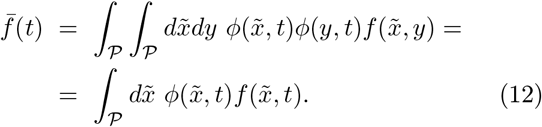

Importantly, Eq.(11), encodes the idea that the marginal fitness of an individual with a given trait, 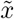, is frequency-dependent, i.e. the fitness associated with a given trait depends crucially on the system’s state, i.e., on the distribution of individuals in phenotypic space.

Even if the derivation of Eq.(10), might look cumbersome, it can be interpreted in a rather straightforward and intuitive way (see the sketch in Fig. 1):

- The first term on the right hand side is a positive probability flow into the *x* state stemming from the probability that an individual with any arbitrary trait 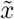is chosen for reproduction and produces a mutated offspring with, precisely, trait *x*.
- The second term describes the fact that, owing to normalization, reproduction events leading to an increase in probability density for any given trait *x*^*′*^ ≠*x* (which occurs with an average rate 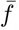) reduce the relative probability of finding individuals in state *x*, i.e. diminish *ϕ*(*x, t*).

### C. Moment equations

Eq. (10) rules the dynamics of the probability density *ϕ*(*x*) and from it one can determine the mean trait

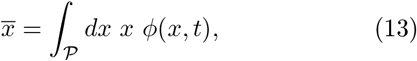

and the central moments

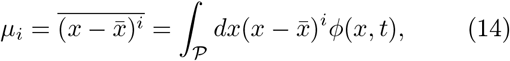

with *i* = 1, 2… and where, in particular, *μ*_1_ = 0, *μ*_2_(*t*) ≡ Σ is the variance, etc. More in general, the mean (population-averaged) value of any possible function *A*(*x*) of the trait *x* is

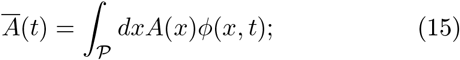

for instance, the mean marginal fitness is 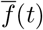. Similarly, the (standard) covariance Σ between any two quantities *A*(*x, t*) and *B*(*x, t*) at time *t* is expressed as

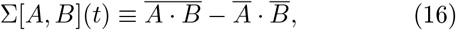

which reduces to the standard variance for *A* = *B*.

From Eq. (10) and the previous definitions, the dynamics of the mean value of an arbitrary function *A*(*x*) is

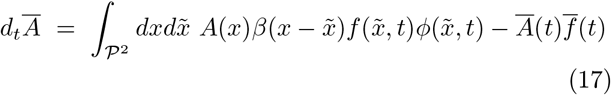

which, if one defines the covariance Σ_*β*_ between two quantities (here *A*(*x*) and *f* (*x*)) across two consecutive generations:

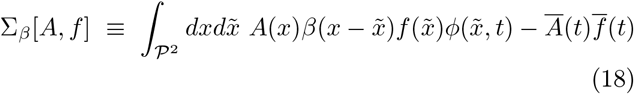

can be written in a very compact form:

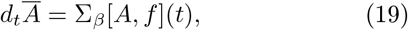

which is nothing but the *Price equation* [17–19, 23]. Observe, in particular, that —while the standard covariance Σ[*A, f*] quantifies the correlation between the quantity *A* and the fitness *f* at a given time— Σ_*β*_[*A, f*] is the covariance between the fitness of the “mother individual” with some trait 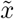 and *A*(*x*) evaluated for the offspring with trait *x* (and the mutation function from 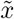 to *x* is included). Thus, Σ_*β*_[*x, f*] stands for the covariance between phenotype and fitness *across a generation*. Note that, in particular, in the absence of variation, i.e. if the mutation amplitude vanishes, both covariances coincide, i.e. 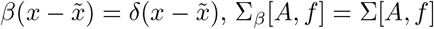.

Summing up, Eq. (19) —the Prize equation— states that if a given function (as for instance, the mean trait value) is positively correlated with the fitness function across a generation its mean-value increases.

### D. Diffusive or small-mutation approximation

Further analytical progress can be made by assuming (as usually done in AD) that the amplitude of mutations 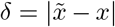 is small, which allows one to perform a series (Kramers-Moyal) expansion of Eq.(10) in powers of *δ* (see SI, Sec.I.C and [104] for details):

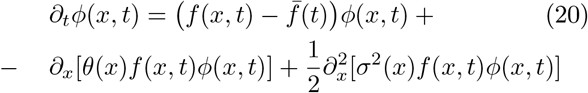

where *θ*(*x*) and *σ*^2^(*x*) are the first two moments of the mutation kernel

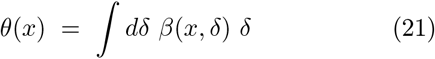

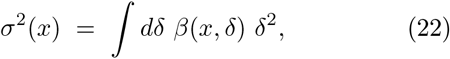

where the first one is referred to as mutation “bias”, the second in the mutation “amplitude”; higher-order terms can be neglected within this approximation, but might be needed in some cases.

Observe that Eq.(20) is a linear superposition of the (upper row) *replicator* dynamics [108], representing selection, and the (lower row) Fokker-Planck type of equation describing mutations as a reaction-diffusion dynamics in phenotypic space. More precisely, the second part is a sort non-linear Fokker-Planck equation or McKean-Vlasov equation [109, 110], as the diffusion function itself depends on the probability-distribution *ϕ*(*x, t*) through the marginal fitness function, Eq.(11), so that it needs to be solved in a self-consistent way.

As a matter of fact, Eq.(20) is actually a version of the celebrated *Crow-Kimura (CK) equation* in population genetics [11, 12] (also called replicator-mutator equation in some contexts [111–114]). Thus, we call Eq.(20) *generalized Crow Kimura equation* (GCK) as it extends the standard CK equation to phenotypic evolution and includes at least two important additional features:

- The fitness function appears in Eq.(20) within the partial derivatives, thus coupling reproduction and mutation.
- Eq.(20) includes generic mutation functions *θ*(*x*) and *σ*^2^(*x*) that, in general, can be trait-dependent rather than constant mutation coefficients.

Let us stress that, within the present small-mutation approximation, given that one can truncate the expansion of the function *β* in power series of *δ*, it is straightforward to recover from Eq.(19), for *A*(*x*) = *x*, after some simple algebra, the more standard form of the Price equation:

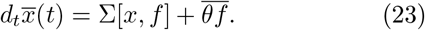

Note also that the contributions of selection and mutation become decoupled in this approximation. In particular, the first term in Eq.(23) encodes *selection* on the mean trait value; i.e. it increases when it correlates positively with fitness increment, while the second term represents the action of biased mutations (so that, in particular, it vanishes if variations are symmetric around *x*) [115]. We could write similar general equations analogous to Eq.(23) for higher moments of the distributions.

For simplicity, let us do so just for the case of constant (trait-independent) mutation rates, for which Eq.(23) becomes

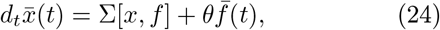

and, similarly, for the trait variance 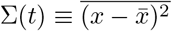 one has:

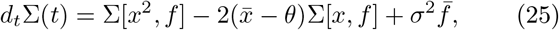

where for simplicity in the notation we have omitted time dependencies.

In general, the equations for higher-order moments form a hierarchy of coupled differential equations. Finding a valid criterion to close such a hierarchy is an open general (“moment-closure”) problem [116]. In what follows, for the sake of completeness and to set the notation and formalism, we first discuss a Gaussian approximation to the closure problem (leading to AD) and then —in forthcoming sections— we introduce extensions accounting for higher-order moments which give rise to a much richer phenomenology.

## III. GAUSSIAN THEORY: RECOVERING ADAPTIVE DYNAMICS

In the classical terms of AD, individuals around the mean phenotype 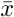 are called “*residents*” and other possible variations of them are called “*mutants*”. As already mentioned, within such a theory, the possible mutations are usually assumed to be small, unbiased, and trait-independent, which greatly simplifies the problem. Under these assumptions, it may suffice to study the dynamics of the mean value 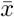 —which coincides with the peak of the underlying distribution— as well as perturbations around it [117].

In other words, to recover the results of AD we consider a *Gaussian approximation* to describe probabilities in phenotypic space, for which one just needs to determine the first two moments —the mean 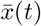 and the variance Σ(*t*)— where the rest are neglected.

More specifically, assuming a general fitness function of the form of Eqs.(2-3), one can expand *f* (*x, y*) in both of its arguments around the mean value 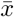:

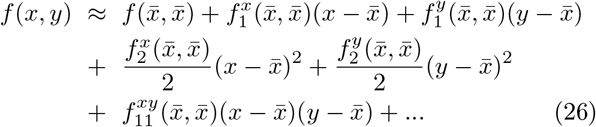

where the subindices indicate the number of derivatives, and the superindices the variables with respect to which the derivatives are taken; e.g.:

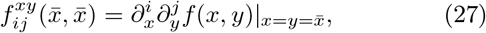

with *i, j* = 1, 2, …

Averaging over the second variable, one obtains the marginal fitness, Eq.(11):

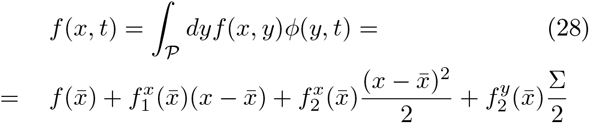

where 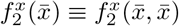 and 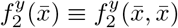, and where both 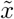 and Σ can in general be time-dependent functions.

Defining 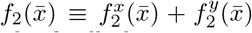, the average fitness, Eq.(12), can be finally be written as:

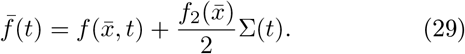

Let us caution that 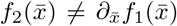 as it also includes the term 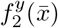; this distinction will be important in what follows.

The previous expressions for *f* (*x*) and 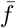, respectively, can be plugged into the Price equation, Eq.(24), to obtain

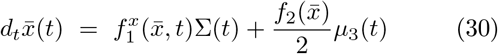

for the case of vanishing bias, *θ* = 0. Moreover, as the phenotypic distribution *ϕ*(*x, t*) has been assumed to be a Gaussian, the third central moment vanishes, i.e. *μ*_3_ = 0, and, hence:

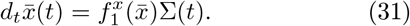

This last equation is known as the “*canonical equation*” in AD and determines the fate of the mean population trait; 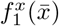 is called “*selection gradient*” and determines the direction of the flow in the in phenotypic space, while the variance of the trait distribution controls the “*speed*” of the evolutionary process [47].

Observe that the possible fixed points, 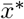, of the previous equation need to be extreme points of the fitness function, i.e. 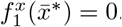. Furthermore, the condition for the stability of the fixed point 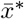 involves the derivative of the fitness gradient with respect to the *mean trait value*:

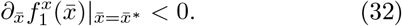

This last condition is called “*convergence stability*” in AD. Let us stress that this condition is different from 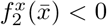, a condition for “evolutionary stability” that we will derive in what follows.

Similarly, from Eq.(25), within the present Gaussian approximation, the variance obeys

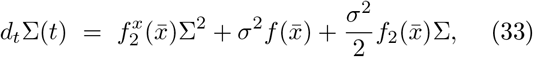

closing set of two equations for the first two central moments of the trait distribution. In particular, the associated steady-state variance is

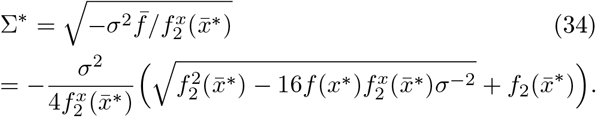

Note that Eq. (34) implies that higher fitness (i.e. larger values of 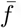) lead to larger steady-state variance. Indeed, larger fitness values means faster reproduction and thus a greater source of mutations and variability. Observe also that the solution of Eq.(34) is a real and stable one only if the second derivative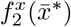 is negative. Therefore, 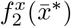 determines whether the variance of the distribution also converges to a steady-state value and thus whether the extreme point 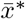 is an *evolutionary stable* state with respect to the introduction of mutants. In other words, an evolutionary stable point needs to be local fitness maximum (see Fig. 2 A).

**FIG. 2.**
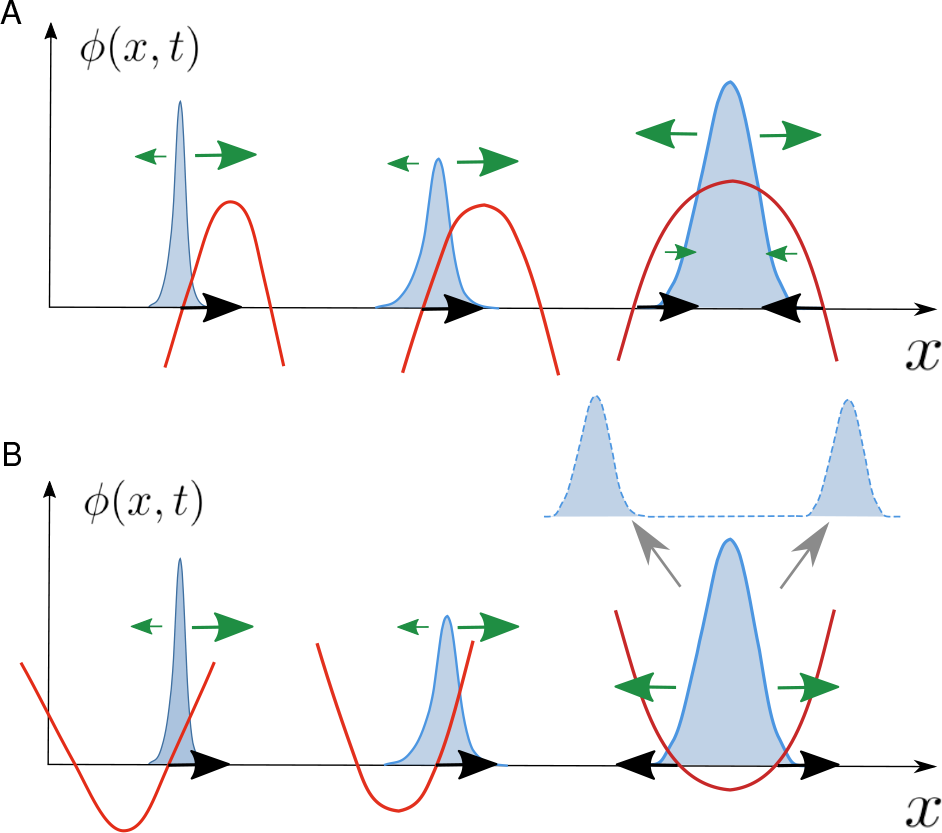
Sketch of typical evolutionary trajectories in adaptive dynamics in the case of (top) a stable monomorphic population and (bottom) evolutionary branching. The distribution of phenotypes is represented at various steps/times of the dynamics, together with the fitness function/landscape (red curves) with changes along with the probability distribution. Green arrows mark the intensity of the selection and the black ones stand for the overall selection gradient. **(A)** Evolutionary stable case: the population climbs the fitness landscape and is attracted to a fitness maximum. **(B)** Evolutionary branching: the population is first attracted to a fitness minimum (evolutionarily unstable) and is then repelled from it. Once the branching occur. Let us remark that, the standard theory of AD is equivalent to considering a Gaussian approximation for small, unbiased, and trait-independent mutations in our generalized framework (though it breakdown after, e.g. evolutionary branching, when the population can not be described as a single mono-modal distribution any more).

On the other hand, if 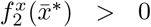, i.e. if it is a minimum, there is no stationary stable solution for the variance and, within the present Gaussian approximation, it just grows unboundedly and 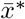 is an attractor for the mean but it is not evolutionarily stable. In this case —recalling that the fitness function is, in general, a dynamic quantity, that changes together with the distribution so that the fitness landscape itself changes during the evolutionary process— the distribution is repelled from the fitness minimum (as illustrated in Fig. 2 B). In this latter case, the evolutionary process (with fixed mean and ever-growing variance) implies that the distribution splits into a bimodal one, with the two peaks progressively diverging from each other, giving rise to the phenomenon of “*evolutionary branching*” [78, 118, 119].

Thus, summing up, under the simplifying assumptions of Gaussian theory, it is possible to explicitly calculate the conditions for the emergence of either evolutionary stable solutions or evolutionary branching, recovering the typical results of AD. In particular, an extreme point of the fitness *x*^*^ is an attractor for the mean trait and — if it is convergent stable, i.e., obeys Eq.(32)— it can lead the population towards two alternative fates:

- an evolutionary stable state emerges when *x*^*^ is a fitness maximum, so that it is a evolutionary stable

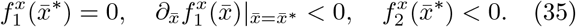
- an evolutionary branching, when 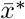 is a fitness minimum, so that there is an evolutionary instability:

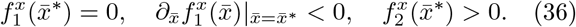

Nevertheless, it is important to emphasize that in order to unveil how the dynamics proceeds beyond a branching point and, more in general, to explore alternative evolutionary phases and patterns in phenotypic space [120–122], it becomes necessary to extend the theory — going beyond the Gaussian approximation— by including higher-order terms in the fitness expansion which may allow, for instance, to describe bimodal or multimodal distributions.

## IV. EXTENDED LANDAU-LIKE THEORY: BEYOND ADAPTIVE DYNAMICS

In order to extend the previous Gaussian theory (i.e. AD) to allow for a description of the evolutionary dynamics even after a branching event —when the variance diverges suggesting that the distribution in phenotypic space can no longer be described as a Gaussian— we now introduce a theory *“a la Landau”* [123, 124] by incorporating higher-order terms in the fitness expansion Eq.(26) (see [50]).

Let us recall that the Landau theory of phase transitions uses a parsimony principle combined with symmetry considerations to write down a general functional —a free-energy functional— including only the most important terms in a perturbative expansion of the relevant field needed to describe key aspects of a given phase transition. For instance, in the classical example of an Ising-like phase transition –describing the spontaneous breaking of an up-down symmetry— one needs to include only even terms and only up to quartic order in a pertubative expansion of the free-energy in powers of the “magnetization” to derive a theory that quantitatively explains the main features of such a transition [123].

Similarly, here we perform an expansion of the “relative fitness”, *F* (*x*), defined as the marginal fitness *f* (*x*) minus its *x*-independent (constant) part:

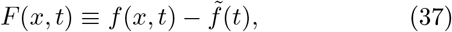

which determines the fitness landscape and is the counterpart of the usual free-energy function.

Before proceeding, let us remark that this relative fitness determines the fitness landscape but, importantly, —much as in statistical mechanics— mutations (which play a role analogous to thermal fluctuations, i.e. temperature) are also crucial in order to determine the shape of the steady-state probability distribution associated with the GCK dynamics, Eq.(20).

Expanding *F* (*x*) in powers of 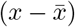 around its mean 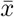 (as in Eq.(26)), one obtains (see SI Sec.III.A):

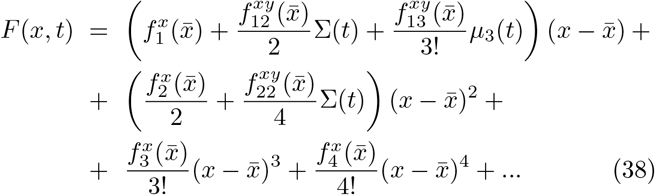

where terms above the 4-th order have been neglected for now.

In order to keep the presentation as simple as possible, let us consider (for now) some simplifying assumptions to *F* (*x*), as usually done in the Landau approach. For example, one can consider the interaction term of the fitness function to be symmetric in its two indices, implying that all the cross-derivative terms of odd order vanish; e.g. 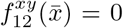. One can also impose the trait distribution *ϕ*(*x, t*) to be symmetric around its mean (at lest in the steady state), which guarantees that all odd central moments, in particular *μ*_3_, vanish at stationarity (note, however, that during the dynamical process the marginal fitness function might not be symmetric even if the fitness function is, owing to asymmetries in the initial fitness distribution.

Under these conditions, the linear term is not changed with respect to the Gaussian theory at stationarity, which implies that 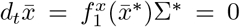, as in the Gaussian approximation; therefore the stationary value, 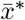, is an extreme point of *F* (*x*), i.e., 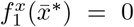. Finally, setting for convenience the stationary mean value to zero, 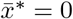, and omitting the dependency on it in the fitness derivatives, the *stationary* relative fitness is:

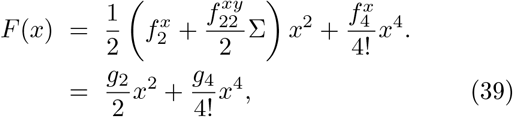

where the functions *g*_2_ and *g*_4_ have been defined in the last line to group the terms proportional to *x*^2^ and *x*^4^, respectively. Let us underscore, once again, the analogy with the Landau theory for the ferromagnetic (Ising) transitions, which has the same formal expansion [123].

The points of vanishing derivative, corresponding either to relative fitness maxima or minima, are the trivial solution, 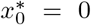, together with and a couple of additional *symmetric* extreme points *x*^*^_1,2_ = *± x*^*^, satisfying:

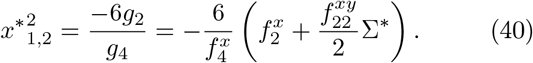

which are real only if *g*_2_ *<* 0 Observe that the location of these two additional fixed points depends on the steady-state variance Σ^*^, something that does not happen in the Gaussian theory.

In order to close the moment hierarchy one needs to determine the steady-state variance using some approximation. For example, one can first assume a bimodal-Gaussian ansatz for the steady-state distribution. However, in spite of the simplification, this is still quite cumbersome to handle analytically, see SI Sec.III B. Thus, we additionally consider the limit of very small mutation amplitudes in which case the bimodal Gaussian becomes the sum of two delta functions,

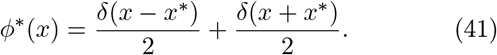

The variance of this distribution is 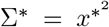 so that plugging this into Eq.(40):

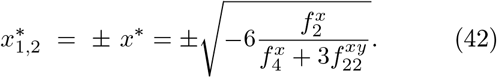

The conditions for these two non-trivial solutions to exist are 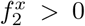 —so that the origin is a fitness minimum (otherwise the Gaussian theory suffices to describe the dynamics)— and 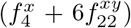, so that there is a positive sign under the square-root and the solutions are real. It is easy to see that 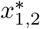 are fitness maxima if 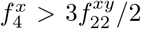, a condition that is expected to be always satisfied when the two points exist and the distribution is bimodal.

Observe that, as already stressed, the sign of 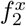 does not fully determine the convexity of *F* (*x*) at the origin (i.e. the overall sign of the terms proportional to *x*^2^ in Eq.(39), i.e. *g*_2_), due to the presence of the additional term 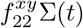, which stems from the interaction kernel as 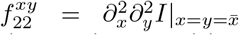 and has the same sign as the interaction: positive for cooperation, negative for competition.

The latter dependence may play a non-trivial role during the course of an evolutionary branching as it will be explicitly illustrated in the next Section. In particular, observe that if the population reaches a fitness minimum and starts branching, the mean-trait value remains fixed in the origin, 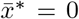 while the variance increases until it possibly reaches a stationary value. This increase in the variance may lead to a progressive change in the coefficient of the quadratic term in Eq.(39), *g*_2_, and hence of the convexity at the origin of the relative fitness function.

To be more specific, one can write down the time derivative of the *g*_2_

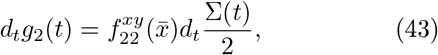

which implies that, e.g., if the variance grows after a branching event, then the fitness barrier separating both of the new attractors can change dynamically, either

- becoming more pronounced if 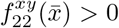 (e.g., for cooperative or mutualistic dominant interactions) so that the two peaks are well separated, or
- becoming flatter if 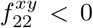 (e.g., for competitive interactions), so that if barrier becomes very flat it may generate a sort of “*neutral bridge*” in between the two coexisting peaks, so that phenotypes with intermediate traits, near *x* = 0 can exist in the steady state even after a branching event [125, 126].

Moreover, a simple calculation including higher non-linearities —such as those appearing in expansions up to 6-th, 8-th, 10-th—-shows that further corrections to the dynamics of *g*_2_ appear. Consequently, the evolutionary dynamics can become much more complex. In particular, it can be easily shown that *g*_2_ can reverse its sign, i.e. the convexity at the origin of the fitness landscape can be inverted after a branching. In this case, a new ecological niche is generated and it can possibly be repopulated, as will be explicitly illustrated by means of a specific example in the next Section. For instance, when after a first branching event, it may happen that each of the two resulting branches converges to a relative fitness minimum, thus leading to a second round of evolutionary branchings and therefore to a total of 4 coexisting sub-populations. If the concavity at the origin has been reverted in the course of the evolutionary process, then the two central branches might eventually converge to the origin colliding and re-populating the empty central niche, resulting in “*evolutionary convergence*” and a final stationary distribution with 3 coexisting sub-populations (see next Section).

Let us remark, that a detailed analytical description of the previously described phenomenology is quite intricate as —in order to allow for the possibility of a series of two branching events— it requires to keep terms up to order *x*^8^ or *x*^10^ in the perturbative expansion of *F* (*x*) and the calculation becomes quite cumbersome [127] (see SI Sec.III C for more details). Thus, in what follows we just move on to present the explicit numerical solution of a specific example where the discussed non-trivial effects such as “neutral bridges”, “cascades of branching events”, and “evolutionary convergence” —all of them beyond the reach of the standard approach of AD— are vividly illustrated and the need of an extended Landau theory becomes also manifest.

## V. GROWTH-COMPETITION MODEL

### A. Model definition

To illuminate the previous theoretical derivations, here we study a simple model that combines characteristics of other models previously introduced in the study of species clustering in phenotypic space [120–122] and in AD [118], respectively. In particular, we consider a one-dimensional phenotypic space, so that individuals are characterized by a single scalar trait, *x*.

Let us first specify the fitness function. The trait value *x* influences the growth rate of individuals, through a growth function *K*(*x*), that can be chosen to be a Gaussian centered at *x* = 0 (plus a constant baseline level). This determines a sort of ecological niche, so that phenotypes close to *x* = 0 are more likely to grow. In addition, individuals with traits *x* and *y* compete with each other with a strength that depends on their trait similarity, i.e., on their distance in phenotypic space, |*x* −*y*|, as specified by certain kernel function *α*(*x*− *y*), that can also be taken to be a Gaussian. Hence, the total fitness of an individual with trait *x*, conditioned to the existence of another individual with trait *y* is specified by

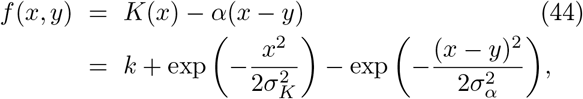

where *k >* 0 is the basal growth rate (warranting the non-negativity of the overall fitness) and the two Gaussians have standard deviations *σ*_*K*_ and *σ*_*α*_, respectively. The model definition is completed by assuming also a Gaussian mutation kernel with zero mean (*θ*0) and *σ* variance. From this

### B. Adaptive dynamics and beyond

Let us first compute the relative fitness function (which depends on the trait distribution) and identify the possible evolutionary phases. Then, we resort to numerical integration of the generalized Crow-Kimura equation, Eq.(20), particularized for the present model. This allows us to gain much insight and visualize the emergent phenomenology. After that we derive some analytical insight by characterizing the steady-state trait distribution using our theoretical framework.

The marginal fitness associated with trait *x* (as defined by Eq.(12), i.e. by integrating the effects of all other possible individuals, *y*, weighted over the trait distribution) reads:

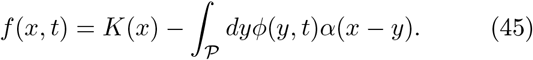

Observe that *K*(*x*) fosters the concentration of the population around the origin and can be seen as an “attracting force” of range *σ*_*K*_, while the competition kernel fosters a kind of “repulsion” between individuals with range *σ*_*α*_. The combined effect of these two forces is illustrated in the two upper leftmost panels of Fig. 3 obtained from numerical integration of the GCK equation for some specific values of the variances and some initial distribution of the population. In particular, the black curves represent the external fitness *K*(*x*), with a maximum at the origin; the green curves stand for the competitive fitness and exhibit a maximum at the point where the population density peaks. Subtracting this second fitness component from the first one, leads to the total fitness function (red curves) which can exhibit a non-symmetric shape. The two upper rows of Fig. 3 illustrate two possible outcomes of the evolutionary dynamics for a monomodal initial distribution, for diverse parameter values.

**FIG. 3.**
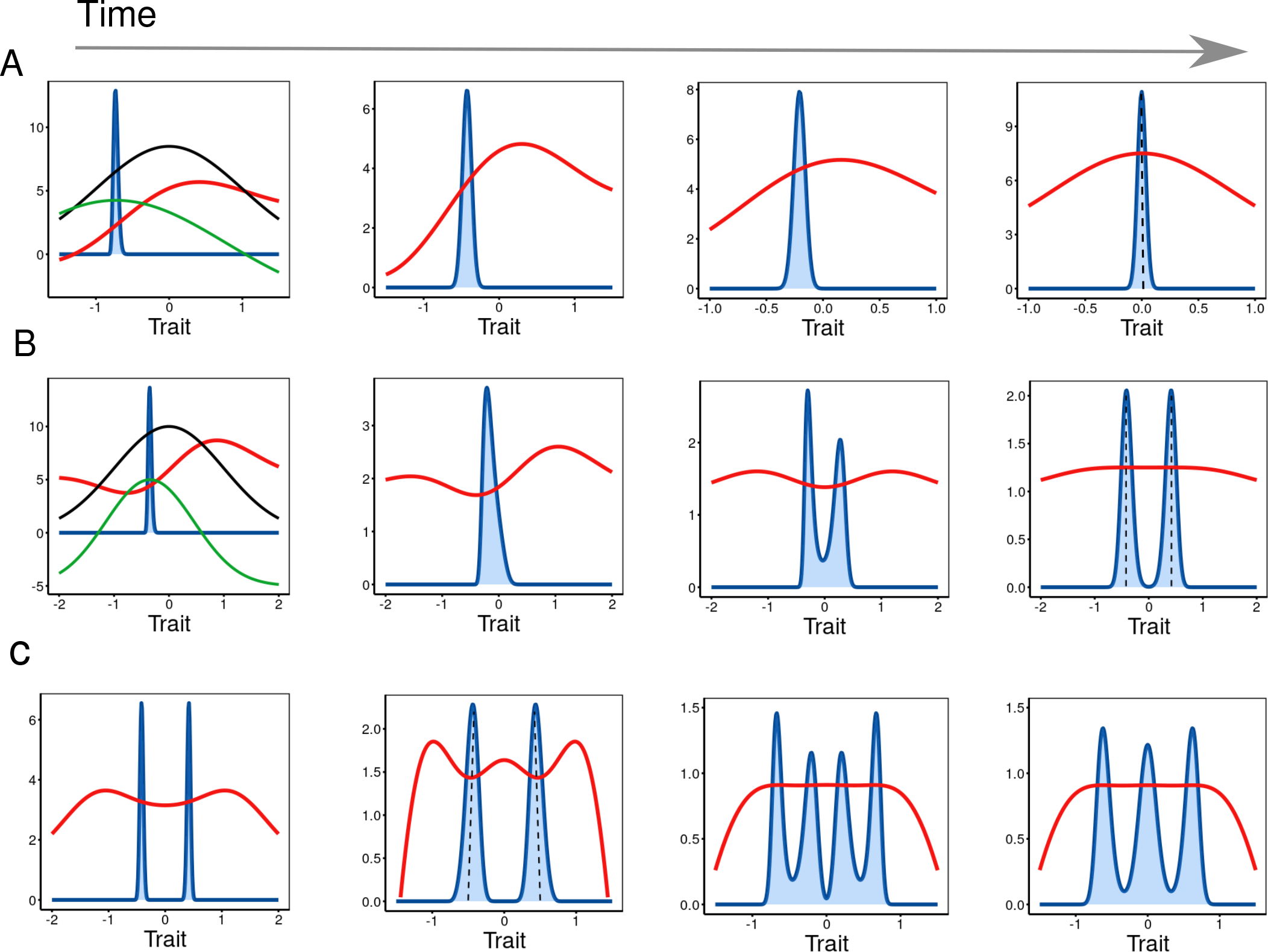
Dynamics of the phenotypic distribution (blue) and fitness landscape (red) for different times as obtained from a numerical integration of the full “generalized Crow-Kimura” (GCK) equation, Eq.(20) in different cases. (A) with no branching, (B) one branching event, and a(C) a series of two consecutive branching and a coalescence event. In the first case (A), the competition kernel (green) is wider than the growth one (black), i.e. *σ*_*α*_ *> σ*_*K*_, producing a fitness landscape with a single maximum (red). The initially peaked distribution moves to the right, in the direction of increasing gradient, thus trying to climb the fitness landscape towards the maximum at *x* = 0, as predicted by theory (dashed vertical line). Observe that the width of the distribution changes across time. In the second case (B), the competition kernel (black) is wider than the growth one, producing a fitness landscape with a two maxima. The population tries to climb the landscape but is trapped in the minimum. To escape and further increase fitness an evolutionary branching happens and the two sub-populations finally reach the the two maxima, as predicted by the theory (dashed lines). Finally, if the competition kernel is way smaller than the growth one (C), after the first evolutionary branching the resulting two sub-populations converge to fitness minima and each one branches again further creating transiently a population with four peaks, that then converges to a 3-peak one, once the two central ones coalesce around *x* = 0. Parameter values: *k* = 1, *σ*_*K*_ = 1 and *σ* = 10^−3^ in all cases. *σ*_*α*_ = 1.2 (A), *σ*_*α*_ = 0.8 (B), *σ*_*α*_ = 0.6 (C). The initial distribution is localized (delta Dirac function) at *x*_0_, with *x*_0_ = −0.8 (A), 0.4 (B,C), respectively.

Before delving into them, let us develop the analytical approach, to be able to make predictions to be compared with numerics.

In order to analyze the dynamics and the possible evolutionary phases, one needs to consider the first derivative of the marginal fitness,

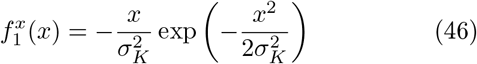

which allows one to determine the direction of the selection gradient as well as to identify the location of its maxima/minima. In particular, *x*^*^ = 0 is always an extreme point, so that the mean value of the distribution converges to 0 along the evolutionary dynamics as predicted by the canonical equation, Eq.(31). It is also convergent stable as

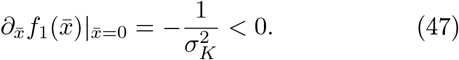

To go beyond standard AD while keeping the calculation as parsimonious as possible, let us expand the relative fitness, 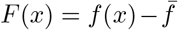, around *x* = 0, including terms up to the 4−th order (as in Eq.(39), above):

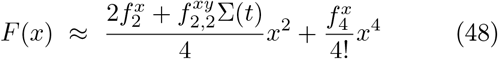

where now we can specify these coefficients

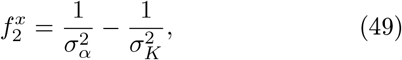

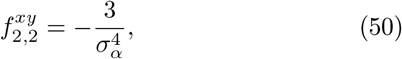

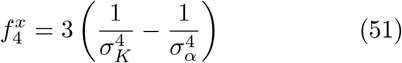

so that 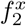 and 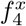 have always opposite signs, while the cross-derivative term, 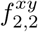, is always negative. Moreover, this last term appears in Eq.(48) multiplied by the variance, so that the full term can be neglected for small mutations. Therefore, from Eq.(49) one readily sees that —at least in an approximate way— the relative fitness has a maximum at the origin 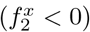 if *σ*_*α*_ ≥ *σ*_*K*_, while the origin is a fitness minimum 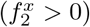 for *σ*_*α*_ *< σ*_*K*_.

Let us discuss these cases separately:

#### (i) No branching

If *σ*_*α*_ ≥ *σ*_*K*_, i.e. the competition kernel has a larger reach than the external fitness, then there is convergence to an evolutionary stable fixed point, i.e. the attractor of the dynamics is a fitness maximum. This is illustrated in Fig. 3 A (obtained from numerical integration of the GCK equation): the leftmost panel shows that when the competition kernel (green curve) is wider than the external-fitness (black curve), the resulting total fitness (red curve) is such that the population climbs the (changing) fitness landscape until the moment in which it stabilizes around the fitness maximum at the origin.

#### (ii) Evolutionary branching

If, on the other hand, *σ*_*α*_ *< σ*_*K*_ the competition kernel has a reach smaller than the external niche; in this case, the attractor is a fitness minimum and the fixed point is evolutionary unstable. This scenario is illustrated in the B series of panels of Fig. 3: after an initial transient the population reaches a fitness minimum, thus leading to a branching event as predicted also by AD. After the branching, the two peaks correspond to extreme points of the overall fitness function. Note that the concavity of the fitness at the origin changes dynamically along the evolutionary process. In particular, after the branching, its becomes flatter and flatter, as qualitatively indicated by our theory (see Eq.(43)) as 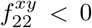, leading to a “neutral bridge”.

#### (iii) Multiple branchings

Finally, by further decreasing *σ*_*α*_ —as explicitly illustrated in the bottom row of Fig. 3— the two peaks shown in the first panel (that may have emerged before in an evolutionary branching event) happen to converge to fitness minima (second panel), so that each of them experiences a second-generation of branching events which —at least, transiently— lead to a distribution with four peaks (third panel); the last (fourth) panel shows that the two central peaks (out of the total 4 transient ones) eventually coalesce together at the origin —repopulating the new niche created after the first branching, as discussed above— and generating a 3-peak steady-state distribution.

By progressively increasing the value of *σ*_*K*_ or diminishing *σ*_*α*_ one can find a cascade of further branching and coalescence events leading to steady-state distributions with a progressively larger number of peaks. Even if results are not shown here, we have performed numerical integration of the GCK and identified steady states with up to 8 peaks (see Fig. 4). This observation can be easily rationalized: *σ*_*K*_ determines the overall width of the available niche space, so the larger *σ*_*K*_ the larger number of diverse phenotypes that can exist. On the other hand *σ*_*α*_ determines the reach of competition, the smaller it is, the closer two consecutive peaks can be, allowing for a more compact packing of “species”. Let us remark that the exact location of the transition lines could in principle be shifted in the case in which 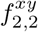 and/or the mutation amplitude, that contribute to *g*_2_ in Eq.(48) are not small.

**FIG. 4.**
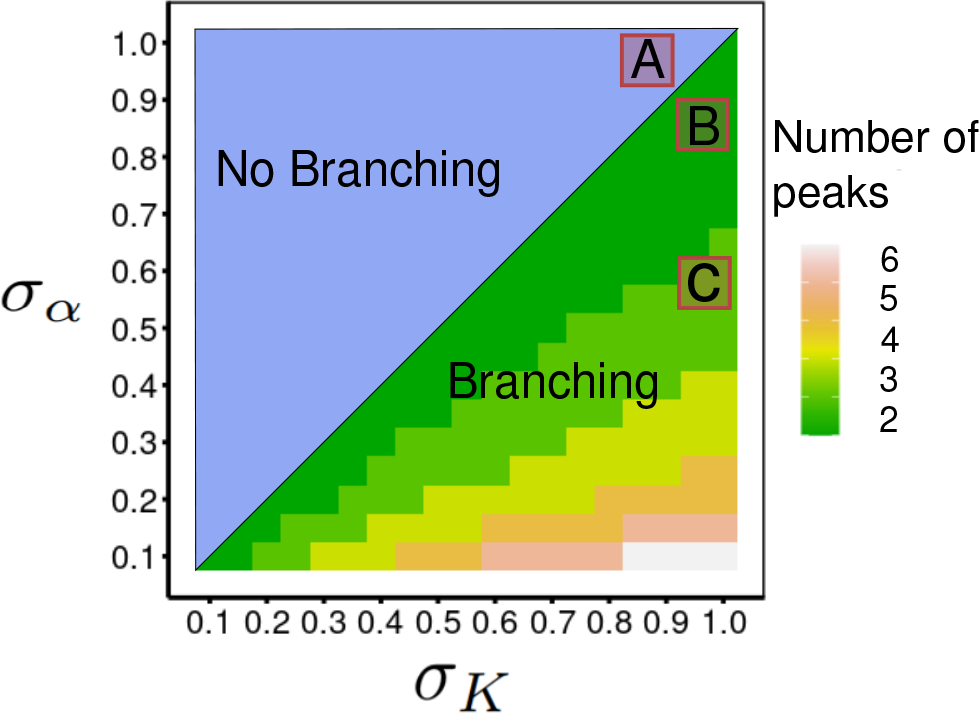
Phase diagram for growth-competition model as a function of the two control parameters *σ*_*α*_ and *σ*_*K*_. Different colors represent the possible phases: uni-modal one with no branching (blue), while the “diversified phase” below the diagonal line (with colors running from green to reddish and white) corresponds to multi-modal distributions (colors codify the number of peaks). The points marked by letters A, B and C identify typical working points for the analyses shown in Fig. 3, with none, one or two branching events, respectively. The diagram is obtained by integrating numerically the generalized Crow-Kimura equation, Eq.(20). The initial condition, as illustrated in Fig. (3), is a delta-Dirac function at *x* = 0.1 in all cases. Other parameter values: *k* = 0.1, *σ* = 10^−3^.

To summarize the previous results, Fig. 4 shows the resulting phase diagram as a function of the two free parameters (*σ*_*α*_ and *σ*_*K*_) controlling the fitness function. In particular, a diversification cascade, consisting in a series of branching events and possible coalescence at the origin may generate steady states with 3, 4, 5, 6, 7 or 8 peaks —as we have computationally verified— though arbitrarily large number of peaks are expected to emerge for sufficiently small values of *σ*_*α*_ and sufficiently large values of *σ*_*K*_.

Finally, it is also noteworthy that by analyzing the time-dependent behavior of the population distribution in situations in which many peaks are expected to occur in the steady state, we observe that in order to reach a final steady state with *n* peaks —starting from a uni-modal distribution at the origin— the probability distribution goes through all the *m < n* possible intermediate phases by a series of evolutionary branching events —i.e. a cascade of dynamical phase transitions— with some coalescence events in between.

### C. Analytical considerations

#### Trait distributions beyond Gaussian theory

Can we explain all the previous phenomenology analytically? Certainly not within the Gaussian approximation. Thus, in order to go analytically beyond AD, we use the generalized Landau framework as introduced in the previous sections to characterize the properties of the stationary trait distribution in the various phases.

When the *σ*_*K*_ ≤ *σ*_*α*_ the population converges to a Gaussian distribution with mean *x* = 0, — coinciding with the fitness maximum— and a non-vanishing variance that can be approximated as (see Eq.(34)):

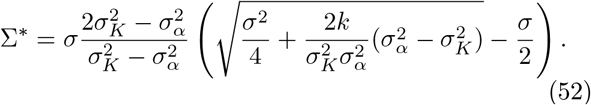

At the transition point, *σ*_*α*_ = *σ*_*K*_ this formula ceases to be valid, as the denominator converges to zero, so that the variance explodes and higher-order terms needs to be considered to estimate the steady-state variance (see SI).

On the other hand, when *σ*_*K*_ *> σ*_*α*_, the population —as already discussed— experiences (at least) one evolutionary branching. To study this phase, it is mandatory to go beyond the quadratic/Gaussian approximation of AD. In particular, as shown above, including up to the 4-th order in the fitness expansion one can explicitly work out the limit of vanishing mutation amplitudes by considering a double delta-function ansatz as specified by Eq.(41). Within such an approximation, the location of the two peaks (as given by Eq.(42)) are:

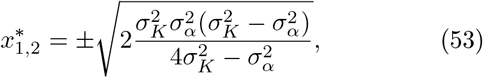

which are real solutions (as *σ*_*K*_ *> σ*_*α*_) and correspond to fitness maxima (see SI).

To increase the precision of the theoretical prediction one could keep terms up to 8-th order in the expansion together with a bimodal Gaussian approximation (i.e. the addition of two symmetric Gaussian distributions) for the stationary distribution. Even if no closed analytical formula is derivable, one can solve numerically equations for the stationary state (see dashed vertical lines in Fig. 3) and compute in particular the mean 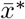 and variance Σ^*^; they match very well the results of integration of the GCK equation for diverse parameter values (see SI Sec.III C for more details).

Fig. 5 summarizes the previous analytical derivations by comparing the stationary variance obtained by numerical integration of the GCK equation (color points) with the theoretical approximations (black line) for different parameter values. Observe that —as illustrated in Fig. 5— by re-scaling both the stationary variance Σ^*^ and *σ*_*α*_ by *σ*_*K*_ all plots collapse to almost the same curve, similarly to what happens for order parameters in second order phase transition [123]. The theoretical prediction compares well with numerics for *σ*_*α*_*/σ*_*K*_ *>* 0.75 while, below such a limit, further higher-order terms are necessary to have a more accurate prediction. In particular, as already discussed, for smaller values of *σ*_*α*_ more peaks appear in the distribution and the bimodal approximation fails and higher-order terms in the Landau expansion are needed.

**FIG. 5.**
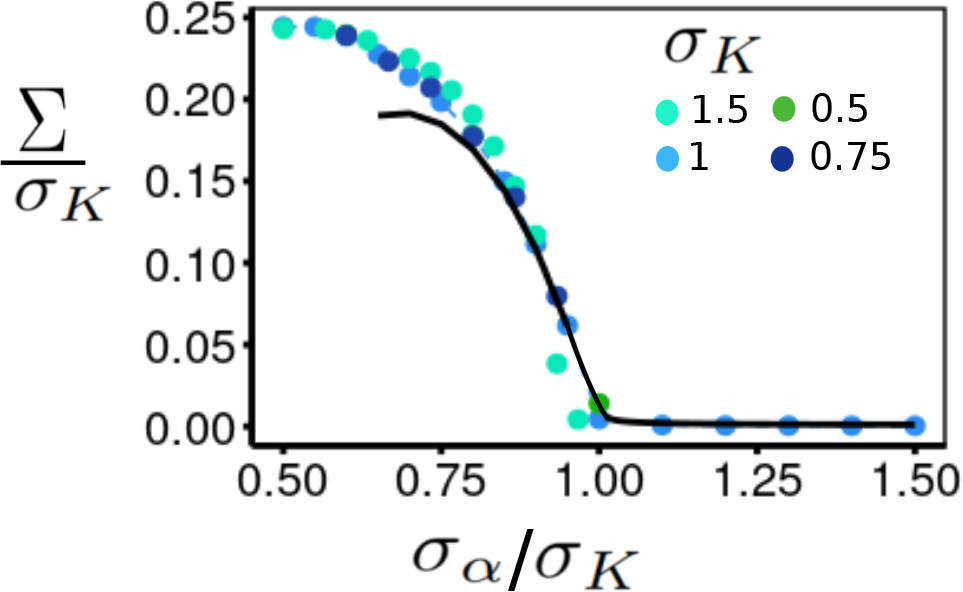
Variance of the steady-state distribution as a function of *σ*_*α*_ (both axes rescaled by *σ*_*K*_), illustrating the analogy with a phase transition. The points represent results from numerical integration of the GCK equation for different values of *σ*_*K*_ (as color-coded in the legend; parameters values as in Fig. 4). Each color represents a fixed value of *σ*_*K*_, while *σ*_*α*_ is changed within the interval [0.5*σ*_*K*_, 1.5*σ*_*K*_]. Observe that thanks to the rescaling by *σ*_*K*_ all different cases approximately collapse onto a single curve. The condition *σ*_*K*_ = *σ*_*α*_ determines the transition from monomorphic populations to evolutionary branching with good accuracy (for the considered mutation variance *σ* = 10^−3^). The black line represents the theoretical predictions obtained from our theory (for *σ*_*α*_ *< σ*_*K*_ we used Eq.(53), corrected with higher terms up to 8-th for *σ*_*α*_ ≤ 0.8*σ*_*K*_, while for *σ*_*α*_ ≥ *σ*_*K*_ we used Eq.(52) corrected by including 6-th-order terms exactly at the transition point *σ*_*K*_ = *σ*_*α*_). The theoretical approximation works relatively well for *σ*_*α*_ *>* 0.75*σ*_*K*_ but deviations are evident for smaller values of *σ*_*α*_, suggesting that higher-order terms are needed in the fitness expansion.

Similarly, we can also estimate the transition lines to the multiple branching phase using numerical evidence. For example, by combining numerical results and the analytical expansion, in Fig. 5 one can see that by further decreasing *σ*_*α*_ the variance reaches a maximum at *σ*_*α*_ = 0.6*σ*_*K*_, below which it continues to decrease. It can be noticed that the maximum coincides with the appearance of the third peak in the stationary distribution, such that the later decrease is due to the emergence of further peaks in between the existing ones, and so with a smaller overall variance.

Summing up, one can conclude, both computationally and analytically, that when the width of the growth kernel, *K*(*x*), is broader than the competition one, i.e. when *σ*_*K*_ *> σ*_*α*_, an evolutionary branching or a series of them emerge in a robust way. Each branching bears strong similarities with continuous phase transitions in statistical physics, with the stationary variance playing the role of an order parameter.

## VI. STOCHASTIC FINITE SIZE FLUCTUATIONS

Demographic effects —stemming from finite-size populations and also known as “genetic drift” in the context of population genetics— are well-known to have a pivotal role in determining the fate of ecological and evolutionary communities. If the population size is relatively small (“population bottleneck” [128, 129]), there might not be variability enough for selection to act upon thus altering the course of evolution. As a matter of fact, the consequences of demographic fluctuations have been extensively studied in the context of population genetics [13, 66], evolutionary game theory [130–134] and AD [80, 88, 98, 135]. In addition to introducing fluctuations around deterministic behavior, demographic fluctuations have been shown to give rise to unexpected phenomenology such as, e.g., evolutionary tunneling, population bottlenecks, and inversion of the direction of selection, to name but a few examples [132, 136, 137].

In order to account for demographic effects in our framework, one needs to move away from the infinite-population-size limit (as developed in Section.II B). In particular, the GCK is exact in the infinite-size limit, so that it needs to be complemented with additional higher-order terms in an 1*/N* expansion.

### A. Computational evidence of demographic effects

Before tackling this problem, let us first explicitly illustrate the difference between the predictions of the previously-derived deterministic theory and the actual stochastic outcome of evolutionary dynamics for finite populations.

For this, we performed computer simulations of the previously defined model master equation, Eq.(4), characterizing the dynamics of the individual-based growth-competition model, by employing an exact Gillespie algorithm (see Appendix A).

Fig. 6 illustrates the results for the evolution of a population of size *N* = 10^3^ and a moderate mutation amplitude, *σ* = 10^−3^, both in the case (A) in which a stationary monomorphic population is predicted at the deterministic level (*σ*_*α*_ *> σ*_*K*_) and in the case (B) in which evolutionary branching is deterministically expected to emerge (*σ*_*α*_ *< σ*_*K*_), as previously summarized in with Fig. 4. The left panels show that the qualitative behavior agrees with the deterministic predictions; however, there is variability across time, i.e. the trajectories are blurred. More specifically, as illustrated in the right panels, the mean value and the variance exhibit stochastic fluctuations around their corresponding steady-state averaged values, which coincide to good approximation with the deterministic expectations (red dashed lines) in all but one of the plots. In the discording case, variance in case (A), there are large asymmetric excursions around certain mean value (blue dashed line) different from the deterministic prediction. Moreover, observe that the mean value in case (B) exhibits fluctuations whose amplitude increases significantly after the population branches out. We will shed light on this in the next Section.

**FIG. 6.**
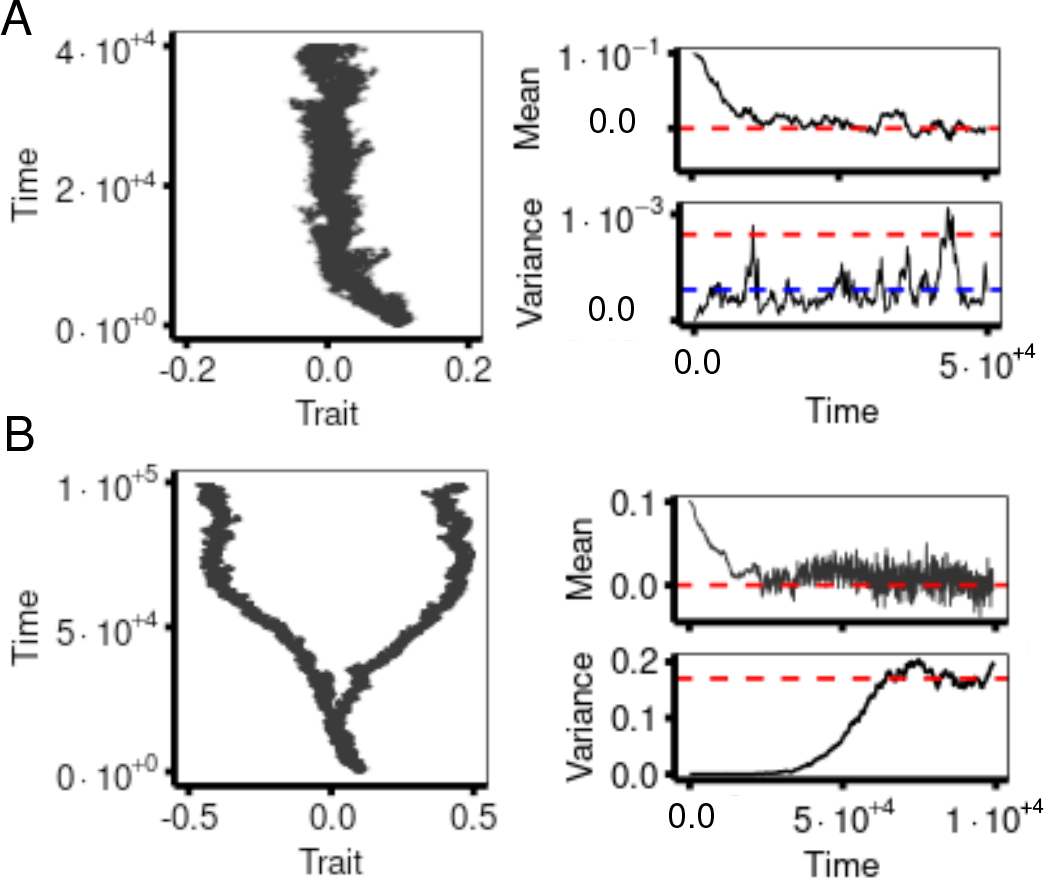
Results of simulations of the individual-based growth-competition model in finite populations. Typical realization of the model dynamics as a function of time for parameter values in the monomorphic/no-branching phase (**Upper panels, A**) and in the phase with one branching event (two peaks) (**Lower panels, B**). Each dot in the left panels corresponds to an individual, thus illustrating the behavior of the population. Instead, the panels to the right show the evolution of the mean population trait (top) and its variance (bottom). The red dashed lines represent the theoretical predictions using Gaussian or Landau theory while the blue one (depicted only for the case in which the Gaussian approximation clearly fails), includes also next-to-leading order stochastic correction, as specified by Eq.(55). Note that in case (A) the variance shows the typical large and asymmetric fluctuations characteristic of multiplicative processes [138, 139]. On the other hand, the fluctuations of the mean in (B) increase significantly after the branching event as suggested by Eqs.(55). The model has been simulated using the individual-based Gillespie algorithm described in detail the Appendix. Parameter values: *N* = 10^3^, *k* = 1, *σ* = 10^−3^, *σ*_*K*_ = 1 and *σ*_*α*_ = 1.2 (A), 0.8 (B).

Similarly, Fig. 7 illustrates the system’s evolution for relatively small population sizes (*N* = 200 and *N* = 1000, respectively) and different mutation amplitude *σ* (*σ* = 10^−3^, *σ* = 2 *×*10^−3^ and *σ* = 5 *×*10^−3^), in the case in which one (upper panels) or more (lower panels) evolutionary branchings are predicted at a deterministic level (i.e. for *σ*_*α*_ = 0.8 and *σ*_*α*_ = 0.6, respectively, with *σ*_*K*_ = 1).

**FIG. 7.**
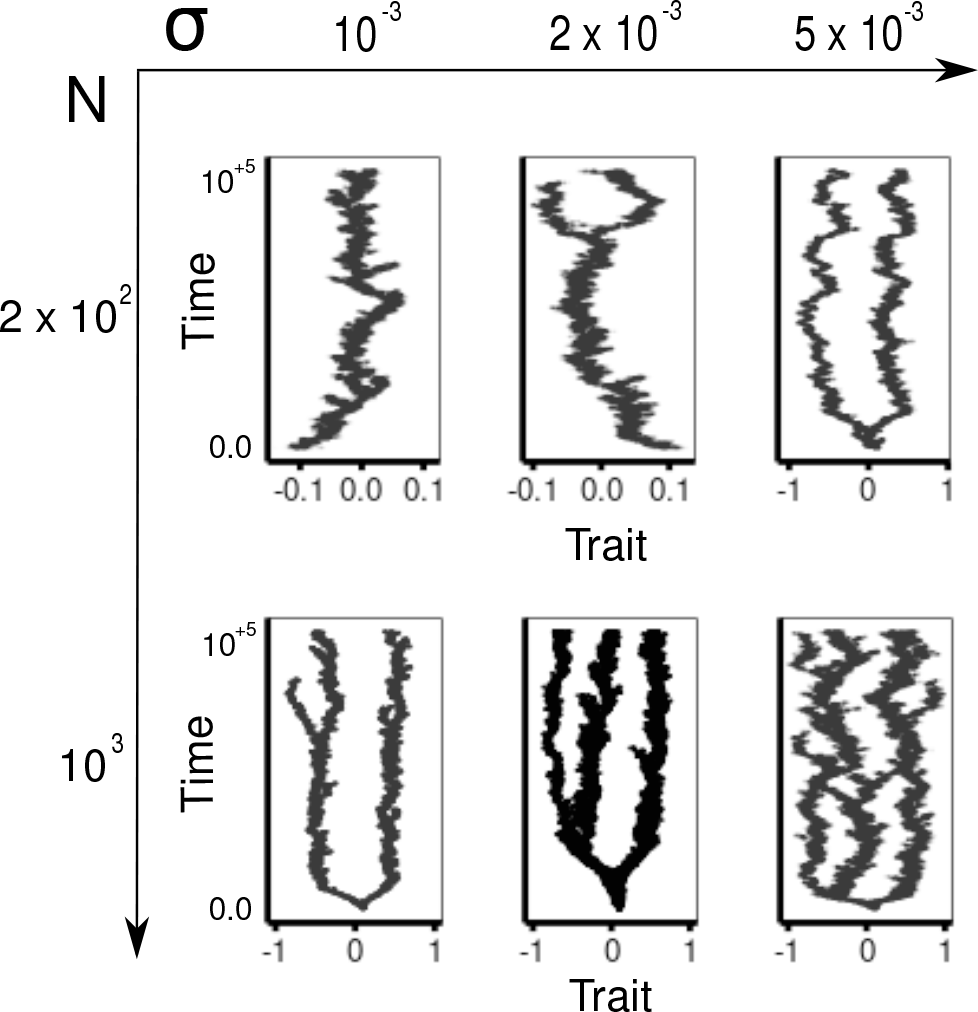
Typical stochastic trajectories for the individual-based growth-competition model illustrating the finite-population effects and their dependence on mutation amplitude *σ* and population size *N*. The evolutionary trajectories of single individuals are plotted (dots) both for the 2-peak phase (*σ*_*K*_ = 1, *σ*_*α*_ = 0.8; upper panels, obtained for *N* = 200) and the 3-peak phase (*σ*_*α*_ = 0.6; lower panels, for *N* = 1000). In the first case, evolutionary branching is not observed (left) for small mutation amplitudes (*σ* = 10^−3^). Branching becomes stochastic, reversible, or “frustrated” (in the sense that one of the branches may become extinct and remaining return to the central position); this occurs for intermediate values of *σ* = 2 *×*10^−3^ (upper central panel) or, alternatively, by slightly increasing *N* (not shown). Finally, branching occurs in a stable and reproducible way (upper right panel) for larger values of the mutation amplitude *σ* = 5 *×*10^−3^ (or larger system sizes). A similar phenomenology can be observed (lower panels) in the 3-peak phase (population size *N* = 10^3^ in this example): for small mutations (lower left panel) the two branches are trapped in their corresponding fitness minima and when additional branchings occur they are frustrated by fluctuations. A stable state with 3 subpopulations occurs for intermediate values of *σ* (lower central panel) or larger values of *N* (not shown) while, finally, a fluctuating 3-peak state appears for larger noise amplitudes.

The top panels (*N* = 200) reveal that for small mutations, *σ* = 10^−3^, there is no stable branching, which implies that the population remains trapped in a fitness minimum. Fluctuations hinder the expected deterministic branching. By increasing *σ* to 2 *×*10^−3^ a tentative but “frustrated” branching appears; i.e. one of the branches becomes extinct and the remaining one moves back to the origin. Finally, for 5 *×*10^−3^, the population is able to generate two branches in a stable way and the two branches seem to keep a constant distance. Thus, as expected, the larger the system size the better the predictions of the deterministic theory are fulfilled.

Similarly, the bottom panels of Fig. 7 show results for a larger population (*N* = 10^3^) in a case (*σ*_*α*_ = 0.6 and *σ*_*K*_ = 1) for which 3 peaks are deterministically expected to emerge. For small mutation amplitudes (*σ* = 10^−3^), much as in the previous case, fluctuations frustrate the emergence of three sub-populations (i.e. the second series of branchings is frustated) and the system remains in a state with just two sub-populations (each of them trapped in a fitness minimum). For larger mutation amplitudes (*σ* = 2 *×*10^−3^) a final state with three sub-populations is reached but —on the contrary of the deterministic predictions (cfr with Fig. 3 C)— the second branching event branching occurs in an asymmetric way (only in the leftmost part in this specific realization). Finally, for even larger amplitudes (*σ* = 5 *×*10^−3^), a complex multi-branching dynamics appears, where the sub-populations branch asymmetrically and wander in phenotypic space, eventually coalescing or going extinct, but keeping three well-separated subpopulations, most of the time.

Summing up, demographic fluctuations have a profound impact on the actual evolutionary/adaptive dynamics that can diverge significantly from the expectations of the deterministic theory. In particular, branching becomes a stochastic phenomenon dependent on the population size and the mutation amplitude and can be “frustrated” if the population is small and/or the mutations are tiny.

### B. Stochastic theory for finite populations

To account for the previously reported deviations from deterministic behavior, in what follows, we present a generalization of our theoretical framework including corrections stemming from finite population sizes. These are derived performing a population size expansion, i.e. a rather standard procedure in the theory of stochastic processes [104] (see SI Sec.IV A).

For simplicity, here we present results just for the case of small, unbiased, and trait-independent mutations (more general cases are discussed in the SI Sec.IV B). Under these restrictions, as shown in detail in the SI, one can derive the following equation for the density in phenotypic space *ρ*(*x, t*) (Eq.(6)), that we call stochastic GCK equation:

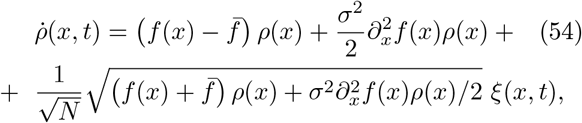

where *ξ* is a delta-correlated, zero-mean, unit-variance Gaussian noise. Note that, in the limit of *N* → ∞, *ρ*(*x, t*) → *ϕ*(*x, t*) and the original GCK equation is recovered. Note that, the square-root functional for in the noise term is the usual one describing demographic noise in birth-death processes and stems from the central limit theorem [104]. Observe also that the stochastic term consists of two contributions:

- The first term, 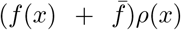, quantifies fluctuations in reproduction and selection (i.e. is associated with the replicator-equation part of the GCK equation).
- The second term, 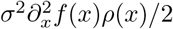 describes fluctuations in mutation events (i.e. is associated with the diffusive part of the GCK equation).

From Eq.(54), it is straightforward to derive a couple of Langevin equations for the mean and the variance, which are the finite-population counterparts of the deterministic equations Eq.(31) and Eq.(33), respectively:

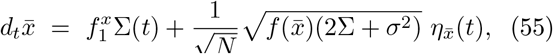

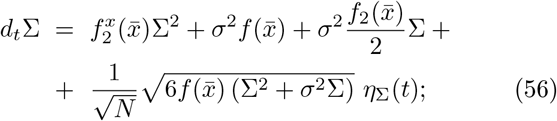

where 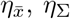 are zero-mean Gaussian white noises (see the SI).

Observe that, importantly, the additional noise term in the equation for the mean-trait value, Eq.(55), grows with the trait variance Σ, explaining why —as illustrated in Fig. 6 B— fluctuations increase significantly after a branching event happens.

On the other hand, the noise in the equation for the variance, Eq.(56), is proportional to the variance itself, i.e. it is a *multiplicative-noise process* [138–140]. This gives rise to a number of remarkable features. First of all, it explains the characteristic asymmetric excursions of the trait variance around its averaged values, observed in Fig. 6 A (second panel in the right column). Second, this multiplicative noise implies that there is an “absorbing state”, meaning that the variance might become “*trapped*” in values very close to zero, from which it is not likely to escape [104, 138–141]. Indeed, as discussed in what follows, this latter effect explains the existence of “frustrated branching” as observed in the left panels of Fig. 7.

#### 1. Interpretation in terms of evolutionary potentials

Let us assume that the mean of the trait distribution has already converged to a given stationary value, 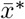, which allows one to have a close Langevin equation for the variance, i.e. Eq.(56) with 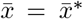. From such a Langevin equation one can readily write its equivalent Fokker-Planck equation and, from it, derive the steady-state probability distribution (see SI Sec.IV C and [104]):

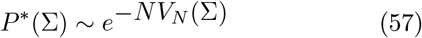

where the effective potential *V*_*N*_ (Σ) is

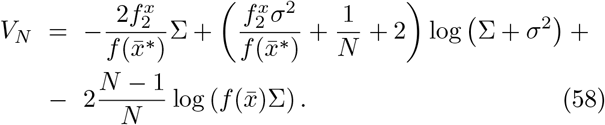

The crucial point is that, for finite values of *N*, the potential *V*_*N*_ (Σ) exhibits a logarithmic singularity at the origin. In particular, in the limit of small mutations, *σ* → 0, the potential becomes:

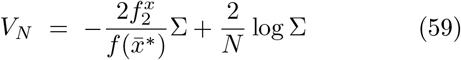

revealing the presence of a negative singularity at small Σ’s (see Fig. 8). The negative singularity at the origin only exists for *σ* = 0, while for small values of *σ* and/or *N*, there is a potential well near the origin; as *σ* and *N* grow the relative weight of the potential well near the origin diminishes and eventually —for sufficiently large sizes and/or mutation amplitudes it disappears (see Fig. 8). This implies that the evolutionary dynamics can become trapped at the potential well around Σ = 0, which induces possible effects absent in the deterministic limit. All this stems from the multiplicative nature of the noise term.

**FIG. 8.**
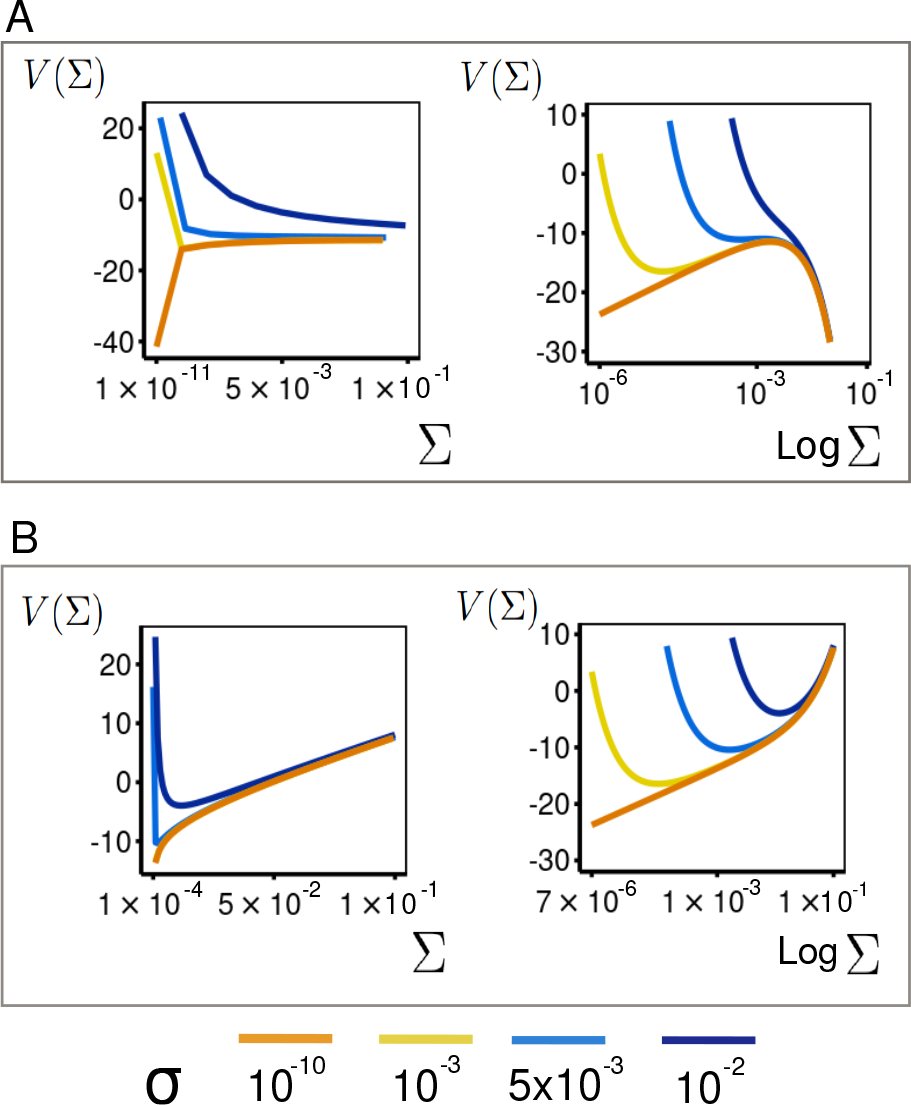
Effective potential, V_N_(Σ) for finite-size populations (N = 200) and variable mutation amplitudes (color coded). The upper panels (A) stand for amplitudes (color coded). The upper panels **(A)** stand for a case in which branching is expected at a deterministic level (i.e. for large populations), while the lower ones **(B)** stand for a case in which a monomorphic population is expected in such a limit. The left panels show potentials in linear scale while the right ones show the same potentials in semi-logarithmic scale. In there are three different regimes depending on the value of *σ* (and *N*). By decreasing the mutation amplitude *σ*, the potential changes from a monotonously decreasing shape (so that the variance is pushed to infinity; blue curve) — so that branching occurs in an almost deterministic way— to another where a meta-stable minimum arises (as can be better appreciated in the semi-logarithmic plot; yellow curve) and branching may occur in a stochastic way if the barrier is overcome. Finally, when *σ*∼ 0 the potential develops a deep well or absorbing state (orange curve) so that the variance remains very small (and the population remains trapped in a fitness minimum). Similarly, in **(B)** the panels reveal a similar transition, but in this case between a situation with a local minimum —characterizing a monomorphic population for relatively large values of *σ* (or *N*)— and a fluctuation-dominated one, in which the minimum moves progressively closer to zero and the potential becomes flatter around it, describing populations with a small variance and large fluctuations around it (see main text). Parameter values: *k* = 1.0, *σ*_*K*_ = 1.0, *σ*_*α*_ = 0.8 (A) and *σ*_*α*_ = 1.2 (B).

#### 2. The case of deterministic branching

The potential *V*_*N*_ (Σ) for the growth-competition model in the case of a relatively small population size *N* = 200 and different values of the mutation amplitude *σ* (as color coded) for the case in which evolutionary branching is deterministically expected to emerge (i.e. the variance grows unboundedly in Gaussian approximation) is plotted in Fig. 8A (both in linear and logarithmic scale). The figure clearly shows that the effective potential can exhibit different shapes (that are better discriminated in the logarithmic plots) depending of the noise amplitude *σ*.

To quantify these possible behaviors, a simple calculation allows one to determine the number of extreme points of the potential as well as their location (see SI Sec.IV C). In particular, one finds that for large values of *N* and *σ*, the potential is monotonously decreasing and that an intermediate local maximum emerges approximately at:

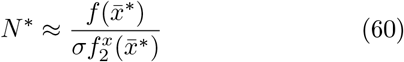

or, in words, when *N* is smaller than a certain (*σ*-dependent) threshold value, *N*^*^, a new “non-deterministic” minimum, together with a local maximum (barrier), appear in the effective potential, describing the dynamics of the variance.

Thus, we can categorize the possible potential shapes as follows.

- *Deterministic branching*. For *N > N*^*^, the potential in Fig. 8A shows a monotonously decreasing curve leading the system to a diverging variance, much as in the deterministic,*N* → ∞, limit. In particular, for large values of Σ, the potential is well approximated by the linear term in Eq.(59), that is proportional to 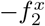, leading to a divergence in variance values (as in the case of deterministic branching, 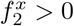). Thus, this phase is dominated by selection and demographic fluctuations are relatively small. At *N* = *N*^*^ the potential develops a saddle point (light blue curve in Fig. 8A).
- *Stochastic branching*. Below the bifurcation point, i.e. for *N* ≲ *N*^*^, the potential exhibits a relative minimum at some value 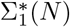 together with a local maximum at 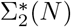 (yellow curve in Fig. 8A). Therefore, a “*drift barrier*” arises between the low variance minimum 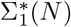 and the large variance regime (that asymptotically leads to divergence and evolutionary branching). Thus, if the variance at the branching point happens to be small, e.g., if the mutation amplitude is small, the population remains trapped in the new minimum. However, possibly fluctuations may drive the population to jump the barrier inducing evolutionary branching stochastically. In this regime finite-size fluctuations, selection and mutation act on similar scales.
- *Frustrated branching*. Finally, if *N* ≪ *N*^*^, the minimum 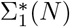 converges to zero [142]. In this limit, the potential converges to Eq.(59) showing a singularity at the origin and a infinitely-high drift barrier between the minimum and the maximum (see orange curves in Fig. 8A). Thus, in this regime, there is a trapping point for small variances, similar to an absorbing state, so that for small sizes and mutation amplitudes *σ*, the probability of stochastically branching becomes arbitrarily close to zero. This regime is thus dominated by drift, and the population cannot diversify.

These three regimes or “phases” are summarized in the phase diagram depicted in Fig. 9 as a function of *σ* and *N* (note the logarithmic scale). The deterministic-branching (light blue) and stochastic-branching (yellow) phases are separated by the critical line *N*^*^(*σ*) (continuous line for computational results and dashed line for the approximated estimation, Eq.(60)), while the “evolutionary absorbing state” or “frustrated branching regime”, emerging for *σ* ≈ 0, is represented as an orange strip to the right-most part of the diagram.

**FIG. 9.**
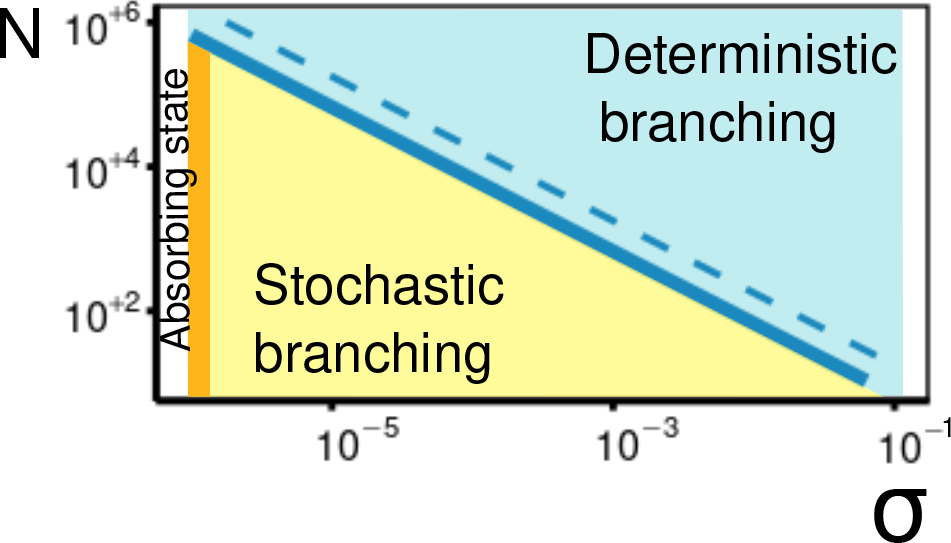
Regimes of branching. Phase diagram of the growth-competition model for arbitrary population sizes, *N* and mutation amplitude *σ*. The transition lines separate the different regimes of branching: *deterministic branching* (blue), *stochastic branching* (yellow) and *frustrated branching or “absorbing state”* (orange). The color code is as in Fig. 8. The blue region corresponds to deterministic branching; the blue continuous line stands for the critical line *N*^*^(*σ*) at which a (saddle-node) bifurcation occurs (as numerically determined in simulations); the blue-dashed line represents the approximation to *N*^*^ given by Eq.(60). By crossing the blue line, the system enters in the stochastic branching regime, with a metastable low-variance state. The probability of reaching the branched state depends on the size of mutations and the height of the barrier.

Observe that these different phases match rather well the computationally observed regimes for the individual-based simulations reported in Fig. 7A (*N* = 200). In particular, in the first plot *N* ≪ *N*^*^ 2 10^3^ and, hence, the population is trapped in the absorbing state; in the second one *N < N*^*^ ≈ 10^3^ and the population tries to stochastically branch but it is ultimately frustrated. Finally, in the rightmost plot, *N* ≲ *N*^*^ 350, the population branches out in a seemingly stable way, even if with fluctuations.

Before concluding, let us emphasize that the fact the demographic effects can hinder population diversification is already known in the context of AD [49, 80, 88]. However, to the best of our knowledge, the phenomenon has not been analyzed in this general mathematical detail nor related to the emergence of absorbing states and new attractors of the evolutionary dynamics [138, 139, 141, 143].

### 3. The case without deterministic branching. Can branching be induced by fluctuations?

Similarly, in the case in which no-branching is expected at a deterministic level (i.e., 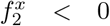), one can see (Fig. 8B) that there is always a single minimum (corresponding to the only solution 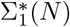 of Eq.(55)), but its associated variance depends on *N* and *σ*. More specifically, for large values of *Nσ* the system behaves almost as in the deterministic case but, as this product decreases, the potential well becomes wider and wider, allowing for larger variability in allowed values of the variance (with progressively smaller mean value, though) i.e. the population develops large intrinsic diversity and temporal variability (see Fig. 8B). In particular, this justifies the findings reported in Fig. 7B for the individual-based model, for which the variance was observed to exhibit anomalous large and asymmetric fluctuations. As a final comment, let us stress that finite-size fluctuations cannot alter the behavior of the effective potential for large values of Σ, so the answer to the question: “can be branching induced by fluctuations?” is “not”, except, maybe, in a transient and reversible way, as illustrated in Fig. 6 A.

## VII. CONCLUSIONS AND DISCUSSION

Historically, microbial evolution has been studied using population genetics. This has been possible owing to the design of high-precision and long-term evolutionary experiments providing access to genetic information and global-fitness measurements of whole (bacterial) populations [55, 144]. Recent studies allowed to generalize classic population-genetics models to rapid or “contemporary” evolution, as observed in actual microbial populations (using e.g. the formalism of statistical physics and concepts such as “fitness travelling waves”) [35]. On the other hand, microbial phenotypic eco-evolution —that was traditionally left aside owing to the difficulties in measuring single-cell traits [145, 146]— has received reinvigorated attention in recent years [84, 147–151] owing to the development of technological advances allowing one to empirically measure traits at an individual-cell level [31, 71, 72] as well as metabolic functions [73–76].

These novel quantitative empirical descriptions of microbial *phenotypic* diversity are crucial for the rapidly-developing field of microbial ecology [105, 152–154] and call for the design of comprehensive eco-evolutionary theoretical frameworks.

This is precisely the problem that aimed to tackle in the present work. In particular, our goal was to analyze populations (with just one species, as a first case study) that can possibly diversify phenotypically into many different “ecotypes” following an eco-evolutionary dynamics (extensions to more complex systems including more than one species and interactions between them is left for future work).

To make progress we have developed a framework that heavily relays on existing ideas in the context of Adaptive Dynamics (AD). We also elaborate upon existing theoretical approaches allowing one to derive “macroscopic” dynamical equations —as those described by e.g. AD— from more “microscopic” dynamical models defined at the level of single individuals and describing the interactions between,, much as done in Statistical Mechanics (some of such previous studies are enumerated and briefly discussed in the Introduction).

First of all, we defined a microscopic individual-based model implementing the fundamental processes of death, reproduction, selection, and mutation. By applying a standard expansion in system (community) size, we first derived a mean-field or deterministic approach that works in an exact way for infinitely-large populations. The resulting dynamics allows us to recover the standard theory of AD by imposing some additional and standard simplifying assumptions (such as, e.g., small mutation amplitudes). However, our description is more general as it allows us to describe not just the evolution of the “mean trait” but the evolutionary dynamics of a whole population in trait space. This dynamcis is encapsulated in a general generalized Crow-Kimura (or replicator-mutator) equation, which remains valid even beyond the previously mentioned approximations. In particular, it extends standard similar equations —previously reported in the literature— in a number of ways as (i) mutations are not necessarily neither small (ii) nor symmetric, and (iii) can be phenotypic-state dependent. Moreover, the coefficients of the GCK depend on the fitness function as both selection and mutation are coupled with reproduction.

To illustrate the generality of our framework, as a first application, in a recent work [155], our group has tackled the problem of the evolution of lag times in bacterial populations exposed periodically to antibiotic stress. For this, we employed a version of the general framework presented here in which the characteristic trait of individuals was the time to wake up from a dormant state (i.e., the “lag time”). Our analyses allowed us to conclude that state-dependent mutations (where the larger the lag the larger the variability of offspring lags) was a necessary ingredient to account for empirical results [155]. The main message from this is that unconventional forms of the mutation kernel can have profound eco-evolutionary consequences in the final distribution of phenotypes, which in turn can have important ecological implications [155].

Similarly, in order o gain further intuition and to put at work the developed formalism, here we have also worked out an explicit example. It consists on a simple individual-based model in which single cells try to occupy and externally imposed niche but also compete among themselves: the “growth-competition model”. A detailed study of this model through the lenses of our theoretical approach allowed us to uncover different eco-evolutionary “phases”, in which the population diversifies in different ways. In particular, the system can either give rise to monomorphic populations (no branching), bimodal populations (on branching event), and, more in general, phases with an arbitrary number of peaks or ecotypes (multiple branching events).

In passing, we illustrated that not all the previous results can be simply explained using the standard (Gaussian) approximation of AD. For instance, to fully describe the bimodal case, one needs to consider corrections up to 4 − *th* order in an expansion of the fitness function around its peak. This resembles what happens in the theory of critical phenomena and phase transitions, where quartic and higher-order non-linearities (in a free-energy expansion) need to be progressively considered to account for more complex (ordered) phases [123].

Last but not least, we have also developed an extension of the framework to account for finite populations and derived an analytical theory that allowed us to describe phenomena such as “stochastic branching”, “frustrated branching”, “evolutionary absorbing states”, etc. — observed, e.g., in simulations of specific model— in a mathematically precise way. This generalized stochastic framework can allow to study other fluctuation-driven evolutionary phenomena such as, e.g., evolutionary tunneling.

Many exciting possibilities open up as follow-ups of the present work that, as already stated, is just a first one of a series.

First, we believe that it is important to use our theory to analyze biologically more structured models, such as consumer-resource ones [83, 147] where different ecological interactions like competition, cross-feeding, environmental fluctuations, etc. maybe simultaneously at work. In particular, environmental fluctuations seem to be the dominant force shaping the statistics of natural microbial communities [105, 153, 156], and little is known about their influence on evolution [157]. To formulate an eco-evolutionary theory able to predict the evolution of metabolic functions in such complex ecological scenarios is a long-term ambitious goal. Similarly, also the influence of spatial effects is emerging as extremely important and will need to be studied [158–161].

Second, related to the previous point, we would like to extend the formalism presented here to systematically account for environmental fluctuations, that can continuously shift the “external niche” to which individuals and communities are exposed and whether population can adapt and possibly diversify following such a change (see [162] for related approaches in AD). In particular, it would be important to analyze the interplay between demographic and environmental fluctuations on the outcome of eco-evolutionary processes.

Third, to further explore the similarities between evolutionary theory and statistical physics we would like to scrutinize whether more complex fitness landscapes, such as rugged ones [5, 6, 163, 164], may lead to chaos [29, 165] and/or replica-symmetry-breaking states [166].

Last but not least, we are presently developing a study of the non-equilibrium properties of evolution (such as entropy production) within our framework, allowing us to quantify “evolutionary irreversibility”, both at a microscopic and macroscopic levels, complementing recent and exciting results [167–170].

Of course, many other items could be added to the previous list of future directions, but we believe it suffices to illustrate the idea that the present work contributes toward the development of a well grounded and fertile theoretical framework to describe phenotypic evolution and complex diversification phenomena, with many potential applications in microbial ecology.

## Appendix A: Simulation algorithm

To simulate exactly and efficiently the microscopic Master Eq.(4) for the growth-competition model, we implemented the exact Gillespie algorithm as follows. Consider N individuals at time *t*_0_:

- The fitness function of all individuals is calculated as the weighted sum over the interaction with all the population:

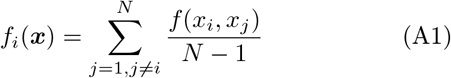

for all *i*. The reproduction probabilities for the different individuals are calculated as 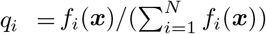.
- A random uniform distributed number *χ* ∈ [0, 1] is generated to calculate the time interval for the next event:

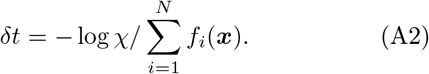
- A random uniform distributed number *η* ∈ [0, 1] is generated and using a standard search algorithm the value of i such that *q*_*i*_ *< η ≤ q*_*i*+1_ is found and the corresponding individual *i* is chosen for reproduction.
- Another individual, say *j*, is randomly chosen with homogeneous probability and is removed from the population. Hence, the j individual is replaced by the offspring of the i individual. Its new trait 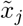 is the mother’s one, *x*_*i*_ plus a random mutation *δ*_*j*_ sampled from a given distribution *β*(*δ*), i.e.:

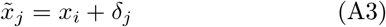
- Time is updated to *t*_0_ = *t*+*δt*. The fitness function of all individuals is updated.
- The process is iterated until a steady state distribution of traits in phenotypic space is reached and the stationary distribution and its moments are measured.

## ACKNOWLEDGMENTS

We acknowledge for financial support (i) the Spanish Ministry and Agencia Estatal de investigación (AEI) through Project of I+D+i Ref. PID2020-113681GB-I00, financed by MICIN/AEI/10.13039/501100011033 and European Regional Development Fund, “A way to make Europe”, and (ii) the Consejería de Conocimiento, Investigación Universidad, Junta de Andalucía and European Regional Development Fund, Project B-FQM-366-UGR20 We also thank J. Grilli, L. Fant, R. Calvo, R. Rubio, J. Knebel, E. Frey, J. Cuesta, J.M. Camacho Mateu, J. Piñero, W. Shoemaker, and L. Peliti for valuable discussions and suggestions. We thank especially R. Corral for a detailed revision of the manuscript.

