## supplementary information for "Statistical mechanics of phenotypic eco-evolution: from adaptive dynamics to complex diversification"

[1]*Departamento de Electromagnetismo y Física de la Materia e Instituto Carlos I de Física Teórica y Computacional. Universidad de Granada. E-18071, Granada, Spain*

**Contents**

**I. From Microsopic to Macroscopic Process: Generalized Moran Process** 1
A. Marginalization 2
B. Mean Field Approximation 4

**II. Kramers-Moyal expansion** 5

**III. Landau-like Theory** 6
A. 4th order expansion 6
B. Bimodal trait distribution 9
C. Higher-order expansion 9

**IV. Finite size fluctuations** 10
A. Derivation from the microscopic process 10
B. Langevin Eqs. for the moments 12
C. Variance analysis 13

**References** 14

### I. FROM MICROSOPIC TO MACROSCOPIC PROCESS: GENERALIZED MORAN PROCESS

In this section we present a microscopic description of evolutionary dynamics, and derive the correspondent mean field equations. Here, the evolutionary model is interpreted as a many-particle Markov jump process, and the mean-field equations are derived by using tools from statistical physics [1, 2]. We consider N *indistinguishable* individuals, or particles, that have a continuous trait $x_i$ in a one-dimensional phenotypic space $\mathcal{P}$. The state of the entire system is represented by the the vector

$$\boldsymbol{x}_N = (x_1, x_2, .x_i, .., x_j, ..x_N), \tag{1}$$
$$dim(\boldsymbol{x}_N) = N; \tag{2}$$

that collects the individual traits $x_i$. Any vector different from $\boldsymbol{x}$ for just one individual trait, say individual j, is indicated as:

$$\widetilde{\boldsymbol{x}}^j_N = (x_1, x_2, ..., x_i, \widetilde{x}_j, ...x_N), \tag{3}$$
$$dim(\widetilde{\boldsymbol{x}}^j_N) = N. \tag{4}$$

In the following paragraphs we study the evolution of probability of a certain configuration $\boldsymbol{x}_N$, $P(\boldsymbol{x}_N)$. The probability, and hence its evolution, satisfies the *symmetry* of individuals indistinguishability that is defined as the fact that the probability is unaffected by any permutation of the elements of the configuration vector $\boldsymbol{x}_N$.

The population evolves by implementing the following stochastic process, that we call a generalized Moran Process:

1 A first individual with trait $x_i$ is selected randomly to reproduce proportionally to its fitness rate $f_i(\boldsymbol{x}_i)$. We restrict our selves to cases where the fitness is composed by a growth rate, $K(x_i)$ and a pairwise interaction $I(x_i, x_k)$:

$$f(x_i, \widetilde{\boldsymbol{x}}^j) \equiv K(x_i) + \frac{1}{N-1}\sum_{k=1, i\neq k}^{N} I(x_i, x_k). \tag{5}$$

2 A second individual with trait $\tilde{x}_j$ is selected randomly to die, i.e. with death rate $d(\tilde{x}_j) = \frac{1}{N-1}$.

3 The second individual dies while the first one reproduces asexually by producing an offspring whose trait will be the parent trait plus a variation, or mutation, $\delta : x_j = x_i + \delta$. The variation is sampled from a probability distribution $\beta(\delta)$ that can in principle depend on the parent trait. Hence, at the end of each event the total number of individuals is conserved.

To translate such process in a stochastic Markov-jump one the 3th needs to be rewritten as a jump. In detail, two jumps are happening: first, the $i$ individual is jumps from position $x_i$ to $x_j$, and the magnitude of the jump is ruled by $\beta(\delta)$. At the same time, the "dying" individual has to jump from $\tilde{x}_j$ to $x_i$, effectively replacing the first individual. This last jump is "non-local" because does not depend on the initial position $\tilde{x}_j$. Such jump is possible thanks individual indistinguishability. By summing up these two jumps, we obtain an effective one from trait $\tilde{x}_j$ to $x_j$, see Fig. 1 in the main text
We can write down a general master equation for the probabilistic dynamics of the the population vector $x$ for the gen. Moran process:

$$\partial_t P(\boldsymbol{x}, t) = \sum_{i=1}^{N} \sum_{j=1, j\neq i}^{N} \int_{\mathcal{P}} d\tilde{x}_j \Big[ W_i(\boldsymbol{x}, \tilde{\boldsymbol{x}}^j) P(\tilde{\boldsymbol{x}}^j, t) - W_i(\tilde{\boldsymbol{x}}^j, \boldsymbol{x}) P(\boldsymbol{x}, t) \Big] \tag{6}$$

where

$$\begin{aligned} W_i(\boldsymbol{x}, \tilde{\boldsymbol{x}}^j) &= f_i(\tilde{\boldsymbol{x}}^j) \;\; \beta(x_j - x_i) \; d(\tilde{x}_j) \\ &= f_i(\tilde{\boldsymbol{x}}^j) \; \beta(x_j - x_i)/(N-1) \end{aligned} \tag{7}$$

is the transition rate to go from a state $\tilde{\boldsymbol{x}}^j = (x_1, .., x_i, .., \tilde{x}_j, .., x_N)$ (with just one coordinate differing from those of $\boldsymbol{x}$) to $\boldsymbol{x} = (x_1, .., x_i, .., x_j, ..x_n)$, and $W_i(\tilde{\boldsymbol{x}}^j, \boldsymbol{x})$ is the rate for the reverse process.

#### A. Marginalization

Now we would like to derive a macroscopic mean field equation for the probability of a trait in the phenotypic space. To do that we formulate a coarse-graining procedure. We introduce the one particle *density* of individual with phenotype $x$ at time $t$:

$$\rho(x, t) = \sum_{i=1}^{N} \frac{\delta(x - x_i)}{N}. \tag{8}$$

Averaging over all possible microscopic configurations one obtains the one-particle probability density

$$\phi(x, t) \equiv \Big\langle \sum_{i=1}^{N} \frac{\delta(x - x_i)}{N} \Big\rangle = \frac{1}{N} \sum_{i=1}^{N} \int_{\mathcal{P}^N} d\mathbf{x} \delta(x - x_i) P(\mathbf{x}, t) = \int_{\mathcal{P}^{N-1}} dx_2 dx_3 ..dx_N P(x, x_2, x_3, .., x_N), \tag{9}$$

where we have renamed the first individual as $x = x_1$,in the second to last passage we used the fact that the particles are indistinguishable(the probability function is symmetric for the exchange of two particles), and in the last one the definition of marginal probability. The same can be written for the n-particle density:

$$\phi^{(n)}(x_1, x_2, .., x_n; t) = \frac{N-n}{N!} \langle \Pi_{j=1}^{n} \sum_{i_j=1}^{N} \delta(x_i - x_{i_j}) \rangle \tag{10}$$

$$= \int_{\mathcal{P}^{N-n}} dx_{n+1} dx_{n+2} ..dx_N P(\boldsymbol{x}, t) = P^{(n)}(x_1, x_2, ..x_n; t). \tag{11}$$

For the moment let us consider just the one particle density Eq.(9) and derive it to respect to time using the Master Eq.(6) to obtain

$$\begin{aligned} \partial_t \phi(x, t) &= \int_{\mathcal{P}^{N-1}} dx_2 ..dx_N \partial_t P(\boldsymbol{x}, t) = \int_{\mathcal{P}^{N-1}} dx_N^{\hat{1}} \sum_{i=1}^{N} \sum_{i\neq j, 1}^{N} \int_{\mathcal{P}} d\tilde{x}_j P(\tilde{\boldsymbol{x}}^j, t) \frac{f_i(\tilde{\boldsymbol{x}}^j, t)}{N-1} \beta(x_j - x_i) + \\ &- \int_{\mathcal{P}^{N-1}} dx_N^{\hat{1}} P(\boldsymbol{x}, t) \sum_{i=1}^{N} \sum_{i\neq j, 1}^{N} \frac{f_i(\boldsymbol{x}, t)}{N-1} = W_{gain} - W_{loss} \end{aligned} \tag{12}$$

Now let us consider the loss contribution (we drop the time dependence):

$$W_{loss} = \int_{\mathcal{P}^{N-1}} dx_N^{\hat{1}} P(\boldsymbol{x},t) \sum_{i=1}^{N} \sum_{i\neq j,1}^{N} \frac{f_i(\boldsymbol{x})}{N-1}, \tag{13}$$

and split the sums in the following way:

$$\sum_{i\geq 1} \sum_{j\geq 1, i\neq j} = \sum_{j>1}^{N} \delta_{i1} + \sum_{i>1}^{N} \sum_{j\geq 1, i\neq j}^{N},$$

to obtain:

$$W_{loss} = \int_{\mathcal{P}^{N-1}} dx_N^{\hat{1}} P(\boldsymbol{x},t) f_1(\boldsymbol{x}) + \int_{\mathcal{P}^{N-1}} dx_N^{\hat{1}} P(\boldsymbol{x},t) \sum_{i>1} f_i(\boldsymbol{x}) = \int_{\mathcal{P}^{N-1}} dx_N^{\hat{1}} [f_1(\boldsymbol{x}) + (N-1) f(x_2, \boldsymbol{x})] P(\boldsymbol{x},t) \tag{14}$$

where, once again, we have used the indistinguishability of individuals Eq.(9).
On the other hand, the gain term is:

$$W_{gain} = \int_{\mathcal{P}^{N-1}} dx_N^{\hat{1}} \sum_{i=1}^{N} \sum_{i\neq j,1}^{N} \int_{\mathcal{P}} d\widetilde{x}_j P(\widetilde{x}^j, t) \frac{f_i(\widetilde{\boldsymbol{x}}^j)}{N-1} \beta(x_j - x_i). \tag{15}$$

Let us split the sums in the following way:

$$\sum_{i\geq 1} \sum_{j\geq 1, i\neq j} = \sum_{j>1}^{N} \delta_{i1} + \sum_{i>1}^{N} \delta_{j1} + \sum_{i>1}^{N} \sum_{j>1, i\neq j}^{N}$$

This is equivalent to say that in the first sum we are considering the case when the particle 1 is reproducing while we are summing over all the possible compatible dying individuals j. In the second term the situation is reversed: individual 1 is dying(i.e. becoming an offspring of j) and we are summing on all the possible j. Finally, the last term considers when individual 1 is not involved in the jump process. Starting with the first term, we integrate over $\widetilde{x}_j$:

$$\sum_{j>1}^{N} \delta_{i1} \int_{\mathcal{P}^N} dx_N^{\hat{1}} d\widetilde{x}_j P(\widetilde{\boldsymbol{x}}^j, t) \frac{f_i(\widetilde{\boldsymbol{x}}^j)}{N-1} \beta(x_j - x_1) = \sum_{j>1} \int_{\mathcal{P}^N} dx_N^{\hat{1}} d\widetilde{x}_j P(\widetilde{\boldsymbol{x}}^j) \frac{f_1(\widetilde{\boldsymbol{x}}^j)}{N-1} \beta(x_j - x_1). \tag{16}$$

$$\tag{17}$$

Now, thanks to individuals indistinguishability each term of the sum gives the same contribution, such that we can fix $j = 2$ without loss of generality:

$$(N-1) \int_{\mathcal{P}^N} dx_N^{\hat{1}} d\widetilde{x}_2 P(\widetilde{\boldsymbol{x}}^2) \frac{f_1(\widetilde{\boldsymbol{x}}^2)}{N-1} \beta(x_2 - x_1) = \int_{\mathcal{P}^N} dx_N^{\hat{1}} d\widetilde{x}_2 P(\widetilde{\boldsymbol{x}}^2) f_1(\widetilde{\boldsymbol{x}}^2) \beta(x_2 - x_1). \tag{18}$$

Next, we can integrate the mutation function using the fact that it is the only term depending on $x_2$ and $\int dx_2 \beta(x_2 - x_1) = 1$, leading to:

$$\int_{\mathcal{P}^N} d\widetilde{x}_N^{\hat{1},2} P(\widetilde{\boldsymbol{x}}^2) f_1(\widetilde{\boldsymbol{x}}^2) \int dx_2 \beta(x_2 - x_1) = \int_{\mathcal{P}^N} d\widetilde{x}_N^{\hat{1},2} P(\widetilde{\boldsymbol{x}}^2) f_1(\widetilde{\boldsymbol{x}}^2). \tag{19}$$

By considering the second term and applying the same procedure of above by fixing $i = 2$ one obtains:

$$\sum_{i>1}^{N} \delta_{j1} \int_{\mathcal{P}^N} dx_N^{\hat{1}} d\widetilde{x}_j P(\widetilde{\boldsymbol{x}}^j) \frac{f_i(\widetilde{\boldsymbol{x}}^j)}{N-1} \beta(x_j - x_i) = \int_{\mathcal{P}^N} dx_N^{\hat{1}} d\widetilde{x}_1 P(\widetilde{\boldsymbol{x}}^1) f_2(\widetilde{\boldsymbol{x}}^1) \beta(x_1 - x_2). \tag{20}$$

Finally, let us consider the last term. Thanks to indistinguishability we can fix $i = 2, j = 3$ by multiplying for a factor $(N-1)(N-2)$ :

$$\sum_{i>1}^{N} \sum_{j>1, i\neq j}^{N} \int_{\mathcal{P}^N} dx_N^{\hat{1}} d\widetilde{x}_j P(\widetilde{\boldsymbol{x}}^j, t) \frac{f_i(\widetilde{\boldsymbol{x}}^j)}{N-1} \beta(x_j - x_i) = (N-2) \int_{\mathcal{P}^N} dx_N^{\hat{1}} d\widetilde{x}_j P(\widetilde{\boldsymbol{x}}^3, t) f_2(\widetilde{\boldsymbol{x}}^3) \beta(x_3 - x_2). \tag{21}$$

Next, we can integrate the mutation function using the fact that it is the only term depending on $x_3$ and $\int dx_3\beta(x_3 - x_2) = 1$

$$(N-2)\int_{\mathcal{P}^N} d\tilde{x}_N^{\hat{1},3} d\tilde{x}_3 P(\tilde{\boldsymbol{x}}^3) f(x_2, \tilde{\boldsymbol{x}}^3) \int dx_3 \beta(x_3 - x_2) = (N-2)\int_{\mathcal{P}^N} d\tilde{x}_N^{\hat{1},3} P(\tilde{\boldsymbol{x}}^3) f_2(\tilde{\boldsymbol{x}}^3) \quad (22)$$

Putting the three terms together:

$$W_{gain} = \int_{\mathcal{P}^N} d\tilde{x}_N^{\hat{1},2} P(\tilde{\boldsymbol{x}}^2) f_1(\tilde{\boldsymbol{x}}^2) + \int_{\mathcal{P}^N} dx_N^{\hat{1}} d\tilde{x}_1 P(\tilde{\boldsymbol{x}}^1) f_2(\tilde{\boldsymbol{x}}^1)\beta(x_1 - x_2) + (N-2)\int_{\mathcal{P}^N} d\tilde{x}_N^{\hat{1},3} P(\tilde{\boldsymbol{x}}^3) f_2(\tilde{\boldsymbol{x}}^3) \quad (23)$$

As a second step, we can consider the explicit form of the fitness function and use the indistinguishability of individuals to simplify more the expression. Namely, the fitness function can be written as the sum over the pairwise fitness over the population:

$$f_i(\boldsymbol{x}) = \sum_{k=1,k\neq i}^{N} \frac{f(x_i, x_k)}{N-1} \quad (24)$$

with:

$$f(x_i, x_k) = K(x_i) + I(x_i, x_k). \quad (25)$$

Let us insert explicitly this expression in the loss term and use individual indistinguishability

$$\begin{aligned} W_{loss} &= \int_{\mathcal{P}^{N-1}} dx_N^{\hat{1}} [f_1(\boldsymbol{x}) + (N-1) f_2(\boldsymbol{x})] P(\boldsymbol{x}, t) \\ &= \frac{1}{N-1}\int_{\mathcal{P}^{N-1}} dx_N^{\hat{1}} \left[\sum_{k=2}^{N} f(x_1, x_k) + (N-1)\sum_{k=1,\neq 2}^{N} f(x_2, x_k)\right] P(\boldsymbol{x}, t) \\ &= \int_{\mathcal{P}} dx_2 P(x_1, x_2) f(x_1, x_2) + (N-1)\int_{\mathcal{P}^2} dx_2 dx_2 P(x_1, x_2, x_3) f(x_2, x_3), \end{aligned} \quad (26)$$

where we have fixed $k = 2$ in the first term and $k = 3$ in the second. Similarly, one can reduce the gain term to:

$$W_{gain} = \int_{\mathcal{P}} dx_2 P(x_1, x_2) f(x_1, x_2) + \int_{\mathcal{P}^2} dx_2 dx_3 \big[P(x_2, x_3) f(x_2, x_3)\beta(x_1 - x_2) + (N-2) f(x_2, x_3) P(x_1, x_2, x_3)\big]. \quad (27)$$

By combining them we obtain the marginalized Master equation:

$$\begin{aligned} \partial_t \phi(x, t) &= W_{gain} - W_{loss} && (28) \\ &= \int_{\mathcal{P}^2} dx_2 dx_3 \big[P(x_2, x_3) f(x_2, x_3)\beta(x_1 - x_2) - f(x_2, x_3) P(x_1, x_2, x_3)\big]. && (29) \end{aligned}$$

### B. Mean Field Approximation

To go further with the anlytic derivation it is necessary to perform a mean field approximation for the N-body probability:

$$P(\boldsymbol{x}, t) = \Pi_{i=1}^{N} \phi(x_i, t), \quad (30)$$

that is expected to be true in the thermodynamic limit $N \to \infty$. By applying such ansatz to the marginalized Master Eq.(28)

$$\partial_t \phi(x, t) = \int_{\mathcal{P}} dx_2 dx_3 \phi(x_2)\beta(x_1 - x_2)\int_{\mathcal{P}} dx_3 \phi(x_3) f(x_2, x_3) - \phi(x, t)\int_{\mathcal{P}} dx_2 \phi(x_2)\int_{\mathcal{P}} dx_3 \phi(x_3) f(x_2, x_3), \quad (31)$$

and by defining the "marginal fitness" as:

$$f(x, t) = \int_{\mathcal{P}} dx d\tilde{x} f(x, \tilde{x})\phi(\tilde{x}, t), \quad (32)$$

one finally obtain the Mean-field mutation-selection Eq.:

$$\partial_t\phi(x,t) = \int_{\mathcal{P}} d\tilde{x} f(\tilde{x},t)\beta(x-\tilde{x})\phi(x,t) - \phi(x,t)\bar{f}(t) \tag{33}$$

with the average fitness defined by

$$\bar{f}(t) = \int_{\mathcal{P}} dx f(x,t)\phi(x,t) = \int_{\mathcal{P}^2} dx d\tilde{x} f(x,\tilde{x})\phi(x,t)\phi(\tilde{x},t). \tag{34}$$

### II. KRAMERS-MOYAL EXPANSION

We are interested in applying a small mutation approximation in the mean-field master Eq. (34), i.e. considering that the offspring trait $x$ is a small deviation from the ancestor, $x = \widetilde{x} + \delta$, $\delta \ll 1$. We consider a general mutation kernel $\beta(x-\widetilde{x};\widetilde{x})$ where we leave the freedom of the dependence on the ancestor trait $\widetilde{x}$. This must be normalized in the jump amplitude $\delta = x - \widetilde{x}$ such that the new trait is still inside the phenotypic space $\mathcal{P}$, that for simplicity we consider as an interval $[p_1, p_2]$

$$\int_{p_1-\widetilde{x}}^{p_2-\widetilde{x}} d(x-\widetilde{x})\beta(x-\widetilde{x};\widetilde{x}) = \int_{p_1-\widetilde{x}}^{p_2-\widetilde{x}} d\delta\beta(\delta;\widetilde{x}) = 1, \tag{35}$$

furthermore, we consider it to be symmetric in the first variable:

$$\beta(\delta, x) = \beta(-\delta, x), \tag{36}$$

and with finite first and second moment:

$$\theta(x) \ = \int_{p_1-\widetilde{x}}^{p_2-\widetilde{x}} d\delta\delta\beta(\delta;x), \qquad \sigma^2(x) = \int_{p_1-\widetilde{x}}^{p_2-\widetilde{x}} d\delta\delta^2\beta(\delta;x). \tag{37}$$

To this aim we rewrite the equation separating the part involving mutations from the rest:

$$\dot{\phi}(x,t) = \Delta_1\phi(x,t) + \Delta_2\phi(x,t) \tag{38}$$

where

$$\Delta_1\phi(x,t) = \int d\widetilde{x}[f(\widetilde{x})\beta(x-\widetilde{x};\widetilde{x})\phi(\widetilde{x},t) - f(x)\beta(x-\widetilde{x};x)\phi(x,t)] \tag{39}$$

and

$$\Delta_2\phi(x) = (f(x) - \bar{f})\phi(x,t) \tag{40}$$

If one consider the first term it is easy to see that it does conserve the probability and has the typical form of a Master equation:

$$\Delta_1\phi(x,t) = \int d\widetilde{x}\Big[W(\widetilde{x},\delta)\phi(\widetilde{x},t) - W(x,\delta)\phi(x,t)\Big] \tag{41}$$

with transition rate

$$W(\widetilde{x},\delta x) = f(\widetilde{x})\beta(\delta;\widetilde{x}). \tag{42}$$

Consider now that this rate can be written as function of the final trait and of the jump amplitude $\widetilde{x} = x - \delta$:

$$W(\widetilde{x},x) = f(\widetilde{x})\beta(x-\widetilde{x};\widetilde{x}) = f(x-\delta)\beta(\delta;x-\delta) := W(t-\delta;\delta). \tag{43}$$

In the same way perform the change of variable in the equation:

$$\Delta_1\phi(x,t) \ = \int_{x-p_2}^{x-p_1} d\delta[W(x-\delta;\delta)\phi(x-\delta) - W(x;\delta)\phi(x)] \tag{44}$$

where the change of integration variable is:

$$\int_{p_1}^{p_2} d\tilde{x} W(\tilde{x}, x)\phi(\tilde{x}, t) \; = \; -\int_{x-p_1}^{x-p_2} d\delta W(x-\delta;\delta)\phi(x-\delta) = \int_{x-p_2}^{x-p1} d\delta W(x-\delta;\delta) \tag{45}$$

Now we can perform the classical taylor expansion in the *first* variable of the rate, assuming $\delta$ small

$$W(x-\delta;\delta) = W(x;\delta) - \delta\partial_x W(x;\delta) + \frac{\delta^2}{2}\partial_x^2 W(x;\delta) + O(\delta^3) \tag{46}$$

and insert it back in the equation ( thanks to the Pawula theorem it sufficient to calculate the first two moments, if they are finite):

$$\Delta_1\phi(x) = -\partial_x m_1(x)\phi(x,t) + \frac{1}{2}\partial_x^2 m_2\phi(x,t) \tag{47}$$

where

$$m_1(x) \; = \; \int_{x-p_2}^{x-p_1} d\delta\delta f(x)\beta(\delta;x) = f(x)\theta(x), \qquad m_2(x) = \int_{x-p_2}^{x-p_1} d\delta\delta^2 f(x)\beta(\delta;x) = \sigma^2(x)f(x) \tag{48}$$

Going back to the full equation we get:

$$\phi(x,t) \; = \; (f(x) - \bar{f})\phi(x,t) - \partial_x\theta(x)f(x)\phi(x,t) + \frac{1}{2}\partial_x^2\sigma^2(x)f(x)\phi(x), \tag{49}$$

that is a generalization the celebrated Crow-Kimura equation [3].

### III. LANDAU-LIKE THEORY

#### A. 4th order expansion

In this section we analyze the fitness function following the parsimonious principle of Landau[4, 5]. In this approach it is fundamental to understand the most important physical properties of the system in order to retain them and neglect the rest. This is achieved by expanding the function representing the properties of the system (the Hamiltonian in statistical physics while the fitness here), and individuate the fundamental terms based on symmetry arguments. Hence, as a first step, we Taylor expand the fitness function around the average value $\bar{x}$ till fourth order to grasp which terms are essential. We start from the basic interaction fitness and expand it in both variables:

$$\begin{aligned} f(x,y) \; \approx \; & f(\bar{x},\bar{x}) + f_1^x(\bar{x},\bar{x})(x-\bar{x}) + f_1^y(\bar{x},\bar{x})(y-\bar{x}) + \frac{1}{2}f_2^x(\bar{x},\bar{x})(x-\bar{x})^2 + \frac{1}{2}f_2^y(\bar{x},\bar{x})(y-\bar{x})^2 \\ + \; & f_{11}^{xy}(\bar{x},\bar{x})(x-\bar{x})(y-\bar{x}) + \frac{1}{3!}f_3^x(\bar{x},\bar{x})(x-\bar{x})^3 + \frac{1}{3!}f_3^y(\bar{x},\bar{x})(y-\bar{x})^3 + \frac{1}{2}f_{21}^{xy}(\bar{x},\bar{x})(x-\bar{x})^2(y-\bar{y}) \\ + \; & \frac{1}{2}f_{12}^{xy}(\bar{x},\bar{x})(x-\bar{x})(y-\bar{y})^2 + \frac{1}{4!}f_4^x(\bar{x},\bar{x})(x-\bar{x})^4 + \frac{1}{4!}f_4^y(\bar{x},\bar{x})(y-\bar{x})^4 + \frac{1}{6}f_{31}^{xy}(\bar{x},\bar{x})(x-\bar{x})^3(y-\bar{x}) \\ + \; & \frac{1}{6}f_{13}^{xy}(\bar{x},\bar{x})(x-\bar{x})(y-\bar{x})^3 + \frac{1}{4}f_{22}^{xy}(\bar{x},\bar{x})(x-\bar{x})^2(y-\bar{y})^2; \end{aligned} \tag{50}$$

where the top indices indicate the variable in which respect the derivative is taken and the bottom one the order of the derivatives, i.e. $f_{22}^{xy}(\bar{x},\bar{x}) = \partial_x^2\partial_y^2 f(x,y)|_{x=y=\bar{x}}$. Once that the interaction fitness expansion has been carried out, by integrating over the trait distribution we obtain the "marginalized" fitness, i.e. the function that appears in the GCK Eq:

$$\begin{aligned} f(x) \; = \; & \int_{\mathcal{P}} dy f(x,y)\phi(y,t) \approx f(\bar{x},\bar{x}) + f_1^x(\bar{x},\bar{x})(x-\bar{x}) + \frac{f_2^x(\bar{x},\bar{x})}{2}(x-\bar{x})^2 + \frac{f_2^y(\bar{x},\bar{x})}{2}\Sigma(t) + \frac{1}{3!}f_3^x(\bar{x},\bar{x})(x-\bar{x})^3 \\ + \; & \frac{1}{3!}f_3^y(\bar{x},\bar{x})\mu_3(t) + \frac{1}{2}f_{12}^{xy}(\bar{x},\bar{x})(x-\bar{x})\Sigma(t) + \frac{1}{4!}f_4^x(\bar{x},\bar{x})(x-\bar{x})^4 + \frac{1}{4!}f_4^y(\bar{x},\bar{x})\mu_4(t) + \frac{1}{6}f_{13}^{xy}(\bar{x},\bar{x})(x-\bar{x})\mu_3(t) \\ + \; & \frac{1}{4}f_{22}^{xy}(\bar{x},\bar{x})(x-\bar{x})^2\Sigma(t); \end{aligned} \tag{51}$$

note that the terms proportional to $(y-\bar{x})$, i.e. $f_1^y, f_{11}^{xy}, f_{21}^{xy}$ and $f_{31}^{xy}$ once integrated are zero because $\int_{\mathcal{P}} dy(y-\bar{x})\phi(y) = (\bar{x}-\bar{x}) = 0$. Finally, by averaging the marginal fitness one obtains the mean one:

$$\begin{aligned}\bar{f} &= \int_{\mathcal{P}} dx f(x)\phi(x,t) = f(\bar{x},\bar{x}) + \frac{f_2^x(\bar{x},\bar{x})}{2}\Sigma(t) + \frac{f_2^y(\bar{x},\bar{x})}{2}\Sigma(t) + \frac{1}{3!}f_3^x(\bar{x},\bar{x})\mu_3(t) + \frac{1}{3!}f_3^y(\bar{x},\bar{x})\mu_3(t) \\ &+ \frac{1}{4!}f_4^x(\bar{x},\bar{x})\mu_4(t) + \frac{1}{4!}f_4^y(\bar{x},\bar{x})\mu_4(t) + \frac{1}{4}f_{22}^{xy}\Sigma(t)^2.\end{aligned} \tag{52}$$

Now, we calculate the relative fitness, i.e. the selection coefficient of the traits. Such a quantity is fundamental because it determines the fittest traits of the GCK Eq.:

$$\begin{aligned}f(x) - \bar{f} &= f_1^x(\bar{x},\bar{x})(x-\bar{x}) + \frac{f_2^x(\bar{x},\bar{x})}{2}[(x-\bar{x})^2 - \Sigma(t)] + \frac{1}{3!}f_3^x(\bar{x},\bar{x})[(x-\bar{x})^3 - \mu_3] + \frac{1}{2}f_{12}^{xy}(x-\bar{x})\Sigma(t) \\ &+ \frac{1}{4!}f_4^x(\bar{x},\bar{x})[(x-\bar{x})^4 - \mu_4] + \frac{1}{6}f_{13}^{xy}(x-\bar{x})\mu_3(t) + \frac{1}{4}f_{22}^{xy}\Sigma(t)((x-\bar{x})^2 - \Sigma).\end{aligned} \tag{53}$$

Here it is fundamental to note that the fitness terms *not* depending on the trait x are canceled out by the mean fitness, and we collect them in the quantity $\widetilde{f}$, while the relevant terms are indicated as $F(x)$. Hence, to determine the fittest traits of the dynamics it is not necessary to study the full fitness function, but just an effective one $F(x)$ that contains the relevant terms and gives the same relative fitness:

$$f(x) = F(x) + \widetilde{f}, \quad f(x) - \bar{f} = F(x) - \bar{F}, \tag{54}$$

where

$$F(x) = f_1^x(\bar{x},\bar{x})(x-\bar{x}) + \frac{f_2^x(\bar{x},\bar{x})}{2}(x-\bar{x})^2 + \frac{1}{3!}f_3^x(\bar{x},\bar{x})(x-\bar{x})^3 + \frac{1}{2}f_{12}^{xy}(x-\bar{x})\Sigma(t)$$
$$+ \frac{1}{6}f_{13}^{xy}(x-\bar{x})\mu_3(t) + \frac{1}{4}f_{22}^{xy}(x-\bar{x})^2\Sigma(t) + \frac{1}{4!}f_4^x(\bar{x},\bar{x})(x-\bar{x})^4 \tag{55}$$
$$\widetilde{f} = f(\bar{x},\bar{x}) + \frac{f_2^y(\bar{x},\bar{x})}{2}\Sigma(t) + \frac{1}{3!}f_3^y(\bar{x},\bar{x})\mu_3(t) + \frac{1}{4!}f_4^y(\bar{x},\bar{x})\mu_4(t). \tag{56}$$

Finally, we can write down the effective fitness function in powers of $(x-\bar{x})$ like the Hamiltonian in statistical physics:

$$F(x) = g_1(t)(x-\bar{x}) + \frac{g_2(t)}{2}(x-\bar{x})^2 + \frac{g_3(t)}{3!}(x-\bar{x})^3 + \frac{g_4(t)}{4!}(x-\bar{x})^4 \tag{57}$$

$$\begin{aligned}g_1(t) &= f_1^x(\bar{x},\bar{x}) + \frac{1}{2}f_{12}^{xy}\Sigma(t) + \frac{1}{6}f_{13}^{xy}\mu_3(t), \quad g_2(t) = f_2^x + \frac{1}{2}f_{22}^{xy}\Sigma(t) \\ g_3(t) &= f_3^x(\bar{x},\bar{x}), \quad g_4(t) = f_4^x(\bar{x},\bar{x}).\end{aligned} \tag{58}$$

First of all, let us note that in principle all the coefficients depend on time, given that the average trait evolves. Second, the first and second order coefficients have some correction terms due to higher-order terms which depend on the trait distribution moments. To better understand this mathematical structure, we follow once more the approach of Landau and search for a symmetry condition valid at *all* times. For example, the trait distribution $\phi(x,t)$ can be assumed to be always symmetric around its mean, imposing$\mu_3(t) = 0$, together with all odd central moments. For simplicity one can also assume the interaction term appearing in the fitness function to be symmetric, leading to vanish all the cross derivative terms of odd order, such as $f_{12}^{xy}(\bar{x})$. In this way, Eq.(57) takes the simpler form:

$$\begin{aligned}F(x) &= f_1^x(\bar{x})(x-\bar{x}) + \left(\frac{f_2^x(\bar{x})}{2} + \frac{f_{22}^{xy}(\bar{x})}{4}\Sigma(t)\right)(x-\bar{x})^2 \\ &+ \frac{f_3^x(\bar{x})}{3!}(x-\bar{x})^3 + \frac{f_4^x(\bar{x})}{4!}(x-\bar{x})^4.\end{aligned} \tag{59}$$

Furthermore, at the stationary state the symmetry condition has further consequences. First, consider the dynamics of the mean trait:

$$d_t\bar{x} = \Sigma\left((x-\bar{x}), f\right) = f_1^x(\bar{x})\Sigma(t) + \frac{f_3^x(\bar{x})}{3!}\mu_4(t), \tag{60}$$

given that the distribution needs to be symmetric around the mean, at the stationary state of the mean $\bar{x}^*$ the third order fitness needs to be zero $f_3^x(\bar{x}^*) = 0$. Hence, this term contributes to the dynamics but vanishes at the stationary state conserving the fact that the stationary mean needs to be an extreme point of the fitness $f_1^x(\bar{x}^*) = 0$. Hence, at the stationary state we can write down an even simpler effective fitness with just two terms :

$$\begin{aligned} F(x) &= \frac{g_2}{2}(x-\bar{x}^*)^2 + \frac{g_4}{4!}(x-\bar{x}^*)^4, && (61)\\ g_2 &= f_2^x(\bar{x}^*) + \frac{1}{2}f_{22}^{xy}(\bar{x}^*)\Sigma, && (62)\\ g_4 &= f_4^x(\bar{x}^*) && (63)\end{aligned}$$

For sake of simplicity we set $\bar{x}^*$=0 without loss of generality. In the case of evolutionary branching, the trait distribution would exhibit two peaks, corresponding to two maxima of the relative fitness Eq(61). To identify them we can take the first derivative and put it to zero:

$$\partial_x F(x) = 0, \tag{64}$$

leading to:

$$x = 0, \qquad \vee \qquad x^2 = {x^*}^2_{1,2} = \frac{-6g_2}{g_4} = -\frac{6}{f_4^x}\left(f_2^x + \frac{f_{22}^{xy}}{2}\Sigma^*\right), \tag{65}$$

showing that the peaks positions would be symmetric, i.e. $x^*_{1,2} = \pm x^*$ but dependent on the variance, and hence on the mutation rate. To solve completely the problem one has to impose a bimodal ansatz on $\phi^*(x)$ and solve both for its mean and variance. This procedure, that ends up to be cumbersome and analytically intractable, is reported in Sec.III B.

Here, we use as a first try a vanishing mutation approximation, assuming that the stationary distribution is a sum of two deltas centered in the extreme points $x^*_{1,2} = \pm x^*$:

$$\phi^*(x) = \frac{\delta(x-x^*)}{2} + \frac{\delta(x+x^*)}{2}. \tag{66}$$

Thanks to this approximation we can determine the variance $\Sigma^* = x^{*2}$ and search for extreme point of the fitness function:

$$\partial_x F(x) = 0, \rightarrow x\left(f_2^x + \frac{f_{22}^{xy}}{2}x^2 + \frac{f_4^x}{3!}x^2\right) = 0 \tag{67}$$

The possible solutions are:

$$x = 0, \quad \vee \quad x = x^*_{1,2} = \pm x^* = \pm\sqrt{-6\frac{f_2^x}{f_4^x + 3f_{22}^{xy}}}, \tag{68}$$

where the first point should be a minimum and the second and third one maxima. The existence condition for $x^*_{1,2}$ are:

$$f_2^x > 0 \qquad \wedge \qquad f_4^x + 3f_{22}^{xy} < 0, \tag{69}$$

or

$$f_2^x < 0 \wedge f_4^x + 3f_{22}^{xy} > 0 \tag{70}$$

On the other hand, the first point needs to be a minimum and hence :

$$\partial_x^2 F|_{x=0} > 0, \rightarrow g_2 = f_2^x + \frac{1}{2}f_{22}^{xy}x^{*2} > 0, \tag{71}$$

by using Eq.(68) this condition turns out to be:

$$f_2^x\left(1 - 6\frac{f_{22}^{xy}}{f_4^x + 3f_{22}^{xy}}\right) > 0 \tag{72}$$

leading to :

$$f_2^x > 0 \qquad \wedge \qquad f_4^x < 3f_{22}^{xy} \tag{73}$$

Then, $x_{1,2}^*$ need to be maxima and hence they should obey:

$$\partial_x^2 F|_{x=\pm x^*} < 0 \quad \rightarrow \quad f_4^x < 0 \tag{74}$$

that is always true for stability.

Summing up, the necessary conditions to have branching are:

$$f_2^x > 0 \quad \wedge \quad f_4^x < 0 \quad \wedge \quad f_4^x < 3f_{22}^{xy}. \tag{75}$$

#### B. Bimodal trait distribution

Here we derive the equation determining the trait distribution in the branching phase using the 4-th order theory and a bimodal ansatz for the stationary distribution:

$$\phi^*(x) = \frac{1}{2}N(-\mu,\omega) + \frac{1}{2}N(\mu,\omega), \tag{76}$$

where $N(\mu,\omega)$ is a normal distribution with mean $\mu$ and std $\omega$. Its central moments are:

$$\begin{aligned} \bar{x} &= \mu_1 = 0, \quad \mu_2 = Var = \omega^2 + \mu^2, \quad \mu_3 = 0 \\ \mu_4 &= 3\omega^4 + 6\mu^2\omega^2 + \mu^4, \quad \mu_5 = 0, \quad \mu_6 = \mu^6 + 14\mu^4\omega^2 + 45\mu^2\omega^4 + 15\omega^6. \end{aligned} \tag{77}$$

From the 4-th order theory the necessary conditions are: $f_2^x(0) > 0$ and $f_4^x(0) < 0$. Note that, in this simplified case, we need to find just two parameters, $\mu$ and $\omega$.

First, by using the extreme condition for $x = \mu$ we obtain a relation between them:

$$\partial_x F(x)|_{x=\mu} = 0 \quad \rightarrow \mu^2 = -6\left(\frac{f_2^x + f_{22}^{xy}\omega^2}{f_4^x + 6f_{22}^{xy}}\right). \tag{78}$$

Second, we consider the stationary equation for the variance $\Sigma$:

$$d_t\Sigma^* = \frac{1}{2}\left(f_2^x + \frac{f_4^{xy}}{2}(\mu_2)\right)(\mu_4 - \mu_2^2) + \frac{f_4^x}{4!}(\mu_6 - \mu_4\mu_2) + \sigma^2\left(f(0) + \frac{f_2^x + f_2^y}{2}\mu_2 + \frac{f_{22}^{xy}}{4}\mu_2^2 + \frac{f_4^x + f_4^y}{4!}\mu_4\right) = 0;$$

by inserting the explicit expression of the moments, Eq.( 77) it becomes:

$$\begin{aligned} &\left(f_2^x + \frac{f_{22}^{xy}}{2}(\omega^2+\mu^2)\right)\omega^2(\omega^2+\mu^2) + \frac{f_4^x}{4!}\left(12\omega^6 + 36\omega^4\mu^2 + 7\mu^4\omega^2\right) \\ &+ \sigma^2\left(f(0) + \frac{f_2^x + f_2^y}{2}(\omega^2+\mu^2) + f_{22}^{xy}\left(\omega^2+\mu^2\right)^2 + \frac{f_4^x + f_4^y}{4!}\left(3\omega^4 + 6\mu^2\omega^2 + \mu^4\right)\right) = 0. \end{aligned} \tag{79}$$

Plugging Eq.(78) into Eq.(79) the problem reduces to solving a cubic equation. Sadly, it is not feasible analytically and one needs to resort to numerical integration. The results are reported in Fig.5 in the main text.

#### C. Higher-order expansion

By following the procedure exposed in Sec. III A one can further and expand the effective fitness at higher orders. For example the expansion up to 6-th order is necessary to calculate the trait variance exactly at the transition point between branching and convergent evolution. On the other hand, order 8-th or higher are necessary to study the deep part of the branching phase. In both of these examples no analytical conclusion can be derived and we relied on numerically solve the involved equations. Here we report briefly the coefficients up to order 8, in the totally symmetric case at the stationary state:

$$F(x) = \frac{g_2}{2}(x-\bar{x})^2 + \frac{g_4}{4!}(x-\bar{x})^4 + \frac{g_6}{6!}(x-\bar{x})^6 + \frac{g_8}{8!}x^8, \tag{80}$$

$$g_2 = f_2^x(\bar{x}) + \frac{f_{22}^{xy}(\bar{x})}{2}\Sigma + \frac{f_{24}^{xy}(\bar{x})}{24}\mu_4 + \frac{f_{26}^{xy}(\bar{x})}{720}\mu_6 \tag{81}$$

$$g_4 = f_4^x(\bar{x}) + \frac{f_{44}^{xy}(\bar{x})}{24}\mu_4 + \frac{f_{42}^{xy}(\bar{x})}{2}\Sigma \tag{82}$$

$$g_6 = f_6^x(\bar{x}) + \frac{f_{62}^{xy}(\bar{x})}{2}\Sigma \tag{83}$$

$$g_8 = f_8^x(\bar{x}), \tag{84}$$

### IV. FINITE SIZE FLUCTUATIONS

#### A. Derivation from the microscopic process

To account for finite-size fluctuations we proceed to derive an equation for $\rho(x,t)$, Eq.(9), that is a random variable describing the density or frequency of individuals with a certain trait x. Note that the average value of the phenotypic density gives the one-particle probability density, and hence recover all previous results. To derive a stochastic equation for $\rho$ we follow the typical procedure used in statistical physics. First, we write down a Master equation in a discrete setting by "coarse-graining" the phenotypic space. Then, by using a Kramers-Moyal expansion [6] in the space of densities under a "local-noise" approximation we derive a Langevin equation for $\rho$. Finally, the continuum limit is performed. We start from the set of $N$ individuals $\boldsymbol{x}_N$ and move to a coarse grained scale where we consider $M$ "species", each one with phenotype $y_i$, abundance $n_i$ and density $\rho_i = n_i/N$ that sum to 1. To go from one scale to another we consider the following clustering, or binning, procedure:

1. Divide the (one-dimensional) phenotypic space $\mathcal{P}$ in M intervals $I_i = ]y_i - \delta_i, y_i + \delta_i]$
2. Individuals with phenotype in the same interval pertain to the same type or "species".
3. Integrate over the individual phenotype and collapse them on the species phenotype. All individuals of the same species have the same fitness, probability, and mutation function. The species fitness is the sum of individual fitness functions.
4. Both the fitness and the mutation function have become discrete functions (elements of a matrix). The mutation function from a species $i$ to $j$ depends on the phenotypic distance of the two.

Hence, starting from the individual probability $P(\boldsymbol{x})$ and its master equation, Eq.(6), first we write down the probability of the species abundances :

$$P(\boldsymbol{n},t) = P(n_1,n_2,..,n_M,t) = \int_{\mathcal{P}^N} d\boldsymbol{x}\delta(x_1-y_{z_1})\delta(x_2-y_{z_2})..\delta(x_N-y_{z_M})P(\boldsymbol{x},t) \tag{85}$$

$$= P(\underbrace{y_1,y_1,..,y_1}_{n_1\text{ times}},\underbrace{y_2,y_2,..,y_2}_{n_2\text{ times}},..,\underbrace{y_M,y_M,..,y_M}_{n_M\text{ times}},t). \tag{86}$$

where the $y_{z_j}$ are the species trait to which they pertain. By taking the time derivative and using the Master equation (6) we get to

$$d_tP(\boldsymbol{n}) = \int_{\mathcal{P}^N} d\boldsymbol{x}_N\delta(x_1-y_{z_1})..\delta(x_N-y_{z_N})\sum_{i,j}[\int_{\mathcal{P}} d\tilde{x}_j f(x_i,\tilde{\boldsymbol{x}}^j)\ \beta(x_j-x_i)d(\tilde{x}_j)P(\tilde{\boldsymbol{x}}^j) - P(\boldsymbol{x})f(x_i,\boldsymbol{x})d(x_j)]$$

Now we apply the coarse-graining instructions

- *Coarse-craining the traits.* One of the delta function, namely the one of $x_i$ acts on the mutation functions:

$$\int dx_i\delta(x_i-y_{z_i})f(x_i,\tilde{\boldsymbol{x}}^j)\ \beta(x_j-x_i)d(\tilde{x}_j)P(\tilde{\boldsymbol{x}}^j) = \beta(y_{z_i}-x_j)f(y_{z_i},\tilde{\boldsymbol{x}}^j)d(\tilde{x}_j)P(\tilde{\boldsymbol{x}}^j) \tag{87}$$

The other $N-1$ deltas act on the fitness and probability giving

$$f(y_{z_i}, \boldsymbol{y}^{z_j,z_{\tilde{j}}})P(\boldsymbol{y}^{z_j,z_{\tilde{j}}}, t) \tag{88}$$

where we have assumed that the new offspring named $x_j$, and the dying individual, $\tilde{x}_j$, pertain respectively to the species $z_j$ and $z_{\tilde{j}}$, and hence the original vector reads:

$$\boldsymbol{y}^{z_j,z_{\tilde{j}}} = (\underbrace{y_1,..,y_1}_{n_1 \text{ times}},..,\underbrace{y_{z_i},..,y_{z_i}}_{n_{z_i} \text{ times}},.., \underbrace{y_{z_{\tilde{j}}},..,y_{z_{\tilde{j}}}}_{n_{z_{\tilde{j}}}+1 \text{ times}},..,\underbrace{y_{z_j},..,y_{z_j}}_{n_{z_j}-1 \text{ times}},..,\underbrace{y_M,..,y_M}_{n_M \text{ times}}) \tag{89}$$

- *From continuous to discrete space.* Second, we rename the indices as $i = z_i, j = z_j$ and $k = z_{\tilde{j}}$, and sum over the possible transitions from state:

$$\tilde{\boldsymbol{n}}^{j,k} = (n_1,..,n_i,..,n_j-1,..,n_k+1,..,n_M) \tag{90}$$

to state:

$$\boldsymbol{n} = (n_1,..,n_i,..,n_j,..,n_z,..,n_M) \tag{91}$$

where a individual of the the species $k$ dies, one of$i$ reproduces by asexually, and the offspring mutates to species $j$. Note that $i,j$ and k can be the same species. Naturally one defines the species fitness and death rate as the sum of all its individuals:

$$f(n_i,\boldsymbol{n}) = \sum_{j\in i} f(x_j,\boldsymbol{x}) = \sum_{j\in i} f(y_i,\boldsymbol{y}) = f(y_i,\boldsymbol{n})n_i, \qquad d(n_i,\boldsymbol{n}) = d(y_i,\boldsymbol{n})n_i. \tag{92}$$

Also, the $\beta$s now are discrete rates $\beta(y_i - y_j) = \beta_{i,j}$.

Finally, adding all the terms:

$$d_t P(\boldsymbol{n},t) = \sum_{i,j,k}[T_i(\boldsymbol{n},\tilde{\boldsymbol{n}}^{j,k})P(\tilde{\boldsymbol{n}}^{j,k},t) - P(\boldsymbol{n},t)T_i(\tilde{\boldsymbol{n}}^{j,k},\boldsymbol{n})] \tag{93}$$

where

$$T_i(\boldsymbol{n},\tilde{\boldsymbol{n}}^{j,k}) = f(y_i,\tilde{\boldsymbol{n}}^{j,k})n_i d(y_k)(n_k+1)\beta_{i,j}. \tag{94}$$

We are now interested in the diffusive approximation in the $N \gg 1$ limit, where we can derive a Langevin equation for the density of i species $\rho_i = \frac{n_i}{N}$ [6]. Consider the following notation for the rates $T_i(\boldsymbol{n},\tilde{\boldsymbol{n}}^{j,k}) = T_i^{j,k}$; to calculate the moments of the expansion let us not that effectively there is just a flux of probability from the k species, who loses an individual, to the j one who obtains an offspring. The probability of the i species does not change and hence we can sum over it. Hence, the first moment of the expansion corresponds to(remember that $d_i = 1/(N-1)$):

$$A_j \;=\; \frac{1}{N}\sum_{i,k}[T_i^{j,k} - T_i^{k,j}] = \frac{1}{N}\sum_{ik}^{M} f_i b_{ij} n_i \frac{n_k}{N-1} - \frac{1}{N}\sum_{ik}^{M} f_i \beta_{ik} n_i \beta_{\frac{n_j}{N-1}}; \tag{95}$$

assuming that $\sum_k n_k = N \approx N-1$, we obtain the deterministic part of the equation:

$$A_j = \sum_{i=1}^{M} \beta_{ij} f_i \rho_i - \bar{f}\rho_j, \tag{96}$$

that is the discretized version of the mean-field equation. In the same way we can calculate the diffusion matrix :

$$B_{jk} \;=\; \frac{1}{N^2}\sum_{i}^{M}[T_i^{j,k} + T_i^{k,j}] = \frac{1}{N}[f_i m_i \beta_{ij} + f_j m_j \beta_{ji} + \sum_{k}^{M} f_k m_k(\beta_{ki} m_j + \beta_{kj} m_i)] \tag{97}$$

$$B_{jj} \;=\; \frac{1}{N^2}\sum_{i,k,k\neq j}^{M}[T_i^{j,k} + T_i^{k,j}] = \frac{1}{N}\sum_i f_i b_{ij}\rho_i + \frac{1}{N}\sum_i f_i \rho_i \rho_j, \tag{98}$$

where we used that $\sum_k \beta_{ik} = 1$. Now, to write down a Langevin equation for $\rho_i$, it would be necessary to diagonalize the matrix B and find the matrix C that follows the relation $C^T C = B$ to obtain:

$$\dot{\rho}_j = A_j + \sum_{k=1}^{M} C_{ik}\xi_k. \tag{99}$$

As clearly expressed in [6] this can be done numerically but not analytically in full generality. Given that our aim here is to derive an analytical expression for such finite size fluctuations in the limit of *infinite* species, it is natural to assume a "local noise approximation", i.e. $B_{jk} = 0$ for $j \neq k$. In this case the Langevin equation is trivially determined as:

$$\dot{\rho}_j = \sum_{i=1}^{M} \beta_{ij} f_i \rho_i - \bar{f}\rho_j + \sqrt{\frac{\sum_i f_i b_{ij}\rho_i + \bar{f}\rho_j}{N}}\xi_j, \tag{100}$$

$$\langle \xi_i(t) \rangle = 0 \tag{101}$$

$$\langle \xi_i(t)\xi_j(t') \rangle = \delta(t-t')\delta_{ij}. \tag{102}$$

By taking the continuum limit, i.e. $M \to \infty$, $\delta \to 0$ we finally obtain the stochastic mutation-selection eq.:

$$\dot{\rho}(x,t) = \int_{\mathcal{P}} d\tilde{x} f(\tilde{x})\rho(\tilde{x})\beta(x-\tilde{x}) - \bar{f}\rho(x) + \sqrt{\frac{\int_{\mathcal{P}} d\tilde{x} f(\tilde{x})\rho(\tilde{x})\beta(x-\tilde{x}) + \bar{f}\rho(x)}{N}}\xi(x,t). \tag{103}$$

Finally, by considering a small-mutation approximation both in the deterministic and stochastic part we end up with the stochastic generalized Crow-Kimura equation:

$$\begin{aligned}
\dot{\rho}(x,t) &= \left(f(x) - \bar{f}\right)\rho(x) - \partial_x \theta f(x)\rho(x) + \frac{1}{2}\partial_x^2 \sigma^2 f(x)\rho(x) \\
&+ \sqrt{\frac{\left(f(x) + \bar{f}\right)\rho(x) - \partial_x \theta f(x)\rho(x) + \frac{1}{2}\partial_x^2\sigma^2 f(x)\rho(x)}{N}}\xi(x,t) \\
\langle \xi(x,t) \rangle &= 0 \\
\langle \xi(x,t)\xi(y,t') \rangle &= \delta(t-t')\delta(x-y).
\end{aligned} \tag{104}$$

In the case of trait-independent and un-biased mutations, one obtains the following simplified stochastic GCK equation:

$$\begin{aligned}
\partial_t \rho(x) &= (f(x) - \bar{f})\rho(x) + \frac{\sigma^2}{2}\partial_x^2 f(x)\rho(x) + \sqrt{\frac{(f(x)+\bar{f})\rho(x) + \frac{\sigma^2}{2}\partial_x^2 f(x)\rho(x)}{N}}\xi(x,t) \\
\langle \xi(x,t) \rangle &= 0 \\
\langle \xi(x,t)\xi(y,t') \rangle &= \delta(x-y)\delta(t-t')
\end{aligned} \tag{105}$$

#### B. Langevin Eqs. for the moments

In this section we will derive a couple of Langevin equation for the trait mean and variance first in a general fashion and then in the context of Gaussian theory. For sake of simplicity in the following we limit ourselves to trait-independent mutations. Consider the mean trait evolution

$$d_t\bar{x}(t) = \int dx(x-\bar{x})\partial_t\rho = f_1\Sigma(t) + \frac{f_2}{2}\mu_3 + \int dx(x-\bar{x})\sqrt{\frac{(f+\bar{f})\rho - \partial_x\theta f\rho + 1/2\partial_x^2\sigma^2 f\rho}{N}}\xi(x,t). \tag{106}$$

where we have used Eq.(104). The stochastic term can be written down as an effective noise in the following form:

$$\eta_{\bar{x}}(t) = \int dx(x-\bar{x})\sqrt{\frac{(f+\bar{f})\rho - \partial_x\theta f\rho + 1/2\partial_x^2\sigma^2 f\rho}{N}}\xi(x,t); \tag{107}$$

$$\langle \eta_{\bar{x}}(t) \rangle = 0 \tag{108}$$

$$\begin{aligned}
\langle \eta_{\bar{x}}(t)\eta_{\bar{x}}(t') \rangle &= \frac{\delta(t-t')}{N}\int dx(x-\bar{x})[(f+\bar{f})\rho - \partial_x\theta f\rho + 1/2\partial_x^2\sigma^2 f\rho] \\
&= \frac{\delta(t-t')}{N}[\overline{(x-\bar{x})^2 f} + \Sigma\bar{f} + 2\overline{(x-\bar{x})\theta f} + \overline{\sigma^2 f}]
\end{aligned} \tag{109}$$

The same can be done for the trait variance $\Sigma$ (where we leave implicit the deterministic part as $G_\Sigma$)

$$d_t\Sigma = G_\Sigma + \int dx(x-\bar{x})^2\sqrt{\frac{(f+\bar{f})\rho - \partial_x\theta f\rho + 1/2\partial_x^2\sigma^2 f\rho}{N}}\xi(x,t) \quad (110)$$

$$\eta_\Sigma(t) = \int dx(x-\bar{x})\sqrt{\frac{(f+\bar{f})\rho - \partial_x\theta f\rho + 1/2\partial_x^2\sigma^2 f\rho}{N}}\xi(x,t) \quad (111)$$

$$\langle\eta_\Sigma(t)\rangle = 0 \quad (112)$$

$$\langle\eta_\Sigma(t)\eta_\Sigma(t')\rangle = \frac{\delta(t-t')}{N}\left(\overline{(x-\bar{x})^4 f} + \mu_4\bar{f} + 4\overline{(x-\bar{x})^3\theta f} + 6\overline{(x-\bar{x})^2\sigma^2 f}\right) \quad (113)$$

The two effective noises are correlated as follows

$$\langle\eta_{\bar{x}}(t)\eta_\Sigma(t')\rangle = \frac{\delta(t-t')}{N}\left(\overline{(x-\bar{x})^3 f} + \mu_3\bar{f} + 3\overline{(x-\bar{x})^2\theta f} + 3\overline{(x-\bar{x})\sigma^2 f}\right). \quad (114)$$

Now by considering un-biased mutations together with the Gaussian approximation, the noise moments reduce to a simpler form:

$$\langle\eta_{\bar{x}}(t)\eta_{\bar{x}}(t')\rangle = \frac{\delta(t-t')}{N}\left(f(\bar{x})(2\Sigma+\sigma) + f_2(2\Sigma+\sigma/2)\right) \quad (115)$$

$$\langle\eta_\Sigma(t)\eta_\Sigma(t')\rangle = \frac{3\delta(t-t')}{N}\left(2f(\bar{x}) + 3f_2\Sigma\right)\left(\Sigma+\sigma^2\Sigma\right) \quad (116)$$

$$\langle\eta_{\bar{x}}(t)\eta_\Sigma(t')\rangle = \frac{3\delta(t-t')}{N}f_1^x(\Sigma+\sigma\Sigma). \quad (117)$$

Finally, we obtain to the couple of simplified Langevin equations that determine the trait distribution in Gaussian theory:

$$d_t\bar{x} = f_1^x\Sigma + \sqrt{\frac{f(\bar{x})(2\Sigma+\sigma) + f_2(2\Sigma+\sigma/2)}{N}}\eta_{\bar{x}}(t) \quad (118)$$

$$d_t\Sigma = f_2^x(\bar{x})\Sigma^2 + \sigma^2 f(\bar{x}) + \frac{\sigma^2}{2}f_2(\bar{x})\Sigma + \sqrt{3\frac{\left(2f(\bar{x}) + 3f_2\Sigma\right)\left(\Sigma+\sigma^2\Sigma\right)}{N}}\eta_\Sigma(t) \quad (119)$$

$$\langle\eta_{\bar{x}}(t)\rangle = 0, \quad \langle\eta_\Sigma(t)\rangle = 0, \quad \langle\eta_{\bar{x}}(t)\eta_{\bar{x}}(t')\rangle = \delta(t-t'), \quad \langle\eta_\Sigma(t)\eta_\Sigma(t')\rangle = \delta(t-t') \quad (120)$$

$$\langle\eta_{\bar{x}}(t)\eta_\Sigma(t')\rangle = \frac{3\delta(t-t')f_1^x(\Sigma+\sigma\Sigma)}{\sqrt{3\left(f(\bar{x})(2\Sigma+\sigma) + f_2(2\Sigma+\sigma/2)\right)\left(2f(\bar{x}) + 3f_2\Sigma\right)\left(\Sigma+\sigma^2\Sigma\right)}} \quad (121)$$

#### C. Variance analysis

To study the stationary behavior of the trait variance, we assume that the mean trait has already converged, enabling us to study the variance independently. Furthermore, for sake of simplicity, we consider just the first fitness term $f(\bar{x})$ in the coefficient of the stochastic noise, obtaining the following simplified equation:

$$d_t\Sigma = f_2^x(\bar{x})\Sigma^2 + \sigma^2 f(\bar{x}) + \frac{1}{2}\sigma^2 f_2(\bar{x})\Sigma + \sqrt{\frac{3f(\bar{x})\left(\Sigma+\sigma^2\Sigma\right)}{N}}\eta_\Sigma(t). \quad (122)$$

Equivalently to Eq.(122), one can consider the following Fokker-Planck equation for the variance probability distribution $P(\Sigma)$( in the Ito discretization scheme):

$$\partial_t P(\Sigma,t) = -\partial_\Sigma[A(\Sigma)P(\Sigma,t) - \partial_\Sigma\frac{B(\Sigma)}{2N}P(\Sigma,t)], \quad (123)$$

with

$$A(\Sigma) = f_2^x(\bar{x})\Sigma^2 + \sigma^2 f(\bar{x}) + \frac{1}{2}\sigma^2 f_2(\bar{x})\Sigma, \quad (124)$$

$$B(\Sigma) = 3f(\bar{x})\left(\Sigma+\sigma^2\Sigma\right). \quad (125)$$

Then, we search for a stationary solution $P^*(\Sigma)$:

$$\partial_t P^*(\Sigma) = 0, \qquad A(\Sigma)P^*(\Sigma) - \partial_\Sigma \frac{B(\Sigma)}{2} P^*(\Sigma) = 0 \tag{126}$$

leading to:

$$P^* = \frac{1}{Z}\exp\left(-V_N(\Sigma)\right) \tag{127}$$

with the potential reading:

$$\begin{aligned} V_N(\Sigma) &= -2N\int d\Sigma \frac{A(\Sigma)}{B(\Sigma)} + \log B(\Sigma) \\ &= \frac{2N}{f(\bar{x}^*)}\left(\left(f_2 - \frac{f_2^x}{2}\right)\sigma^2 + f(\bar{x}^*)\right)\log\left(\Sigma+\sigma^2\right) - 2N\log\Sigma - \frac{2Nf_2^x}{f(\bar{x}^*)}\Sigma - \log\left(f(\bar{x})\left(\Sigma^2+\Sigma\sigma^2\right)\right), \end{aligned} \tag{128}$$

where in the main text for simplicity we have assumed $\left(f_2 - \frac{f_2^x}{2}\right) \approx f_2^x/2$ in the first term. Now, we search for the extreme point of the potential, $\partial_\Sigma V_N(\Sigma^*) = 0$, which leads to the following solution:

$$\Sigma^*_{1,2}(N) = \frac{\sigma^2 b_N}{2f_2^x(x^*)a_N}\left(\pm\sqrt{\sigma^2 f_2^2(x^*) - 4f(x^*)f_2^x(x^*)a_N c_N/b_N} - f_2(x^*)\right) \tag{129}$$

with the size-dependent coefficients

$$a_N = \left(1 - \frac{27f_2}{Nf_2^x}\right), \quad b_N = \left(1 - \frac{3}{Nf_2\sigma^2}\left(2f(\bar{x}) - 3f_2\sigma^2\right)\right), \quad c_N = \left(2 - \frac{3}{N}\right). \tag{130}$$

Depending on the determinant sign $\Delta_N$, Eq.(129) has zero $\Delta_N < 0$, one $\Delta_N$ or two $\Delta_N > 0$ reals solutions, that correspond to the three regimes reported in the main text. The condition of $\Delta_N > 0$ can be translated into a condition for the size N to be smaller than a critical one $N^*$:

$$N < N^* = \frac{f(\bar{x}^*)}{\sigma}\left(\frac{8\sqrt{f(\bar{x}^*)} + 2\sigma\sqrt{f_2^x}}{\sqrt{f_2^x}(16f(\bar{x}^*) - f_2^x\sigma^2)}\right) \approx \frac{f(\bar{x}^*)}{f_2^x\sigma} \tag{131}$$


[1] H. Spohn, *Large Scale Dynamics of Interacting Particles*, Theoretical and Mathematical Physics (Springer Berlin Heidelberg, 2012), ISBN 9783642843716, URL https://books.google.es/books?id=81joCAAAQBAJ.

[2] M.Bauer, J.Knebel, M.Lechner, P.Pickl, and E.Frey, elife **6** (2017), URL https://elifesciences.org/articles/25773.

[3] M. Kimura, Journal of Applied Probability **1**, 177 (1964).

[4] M. Kardar, *Statistical Physics of Fields* (Cambridge University Press, 2007).

[5] J. Binney, N. Dowrick, A. Fisher, and M. Newman (Oxford, 1992).

[6] A. Traulsen, J. C. Claussen, and C. Hauert, Phys. Rev. E **85**, 041901 (2012), URL https://link.aps.org/doi/10.1103/PhysRevE.85.041901.
